# RapA opens the RNA polymerase clamp to disrupt post-termination complexes and prevent cytotoxic R-loop formation

**DOI:** 10.1101/2024.09.28.614012

**Authors:** Joshua J. Brewer, Koe Inlow, Rachel A. Mooney, Barbara Bosch, Paul Dominic B. Olinares, Leandro Pimentel Marcelino, Brian T. Chait, Robert Landick, Jeff Gelles, Elizabeth A. Campbell, Seth A. Darst

## Abstract

Following transcript release during intrinsic termination, *Escherichia coli* RNA polymerase (RNAP) often remains associated with DNA in a post-termination complex (PTC). RNAPs in PTCs are removed from the DNA by the Swi2/Snf2 ATPase RapA. Here, we determined PTC structures on negatively-supercoiled DNA as well as of RapA engaged to dislodge the PTC. We found that core RNAP in the PTC can unwind DNA and initiate RNA synthesis but is prone to producing R-loops. We show that RapA helps control cytotoxic R-loop formation *in vivo*, likely by disrupting PTCs. Nucleotide binding to RapA triggers a conformational change that opens the RNAP clamp, allowing DNA in the RNAP cleft to reanneal and dissociate. We suggest that analagous ATPases acting on PTCs to suppress transcriptional noise and R-loop formation may be widespread. These results hold significance for the bacterial transcription cycle and highlight a role for RapA in maintaining genome stability.

---

RNA-polymerase (RNAP) is the central enzyme of transcription across all domains of life. In the canonical bacterial transcription cycle, the catalytic core RNAP (E; subunit composition α_2_ββ’ω) combines with a promoter-specificity σ factor to form the holoenzyme (Eσ^70^) capable of initiating transcription from specific promoter sequences ^1^. Once transcription initiates and the RNAP becomes committed to elongating the nascent RNA chain, the σ subunit generally releases (although not always) ^2^ and core RNAP completes the elongation and termination phases of the transcription cycle. Although the elongation phase is highly processive, during intrinsic termination the completed RNA transcript is rapidly released from the complex ^3^, followed in the canonical scheme by release of RNAP from the DNA template.

Recent single-molecule investigations have shown that the canonical bacterial transcription cycle must be reevaluated ^4,5^. In these studies, core RNAP was observed to remain associated with the template DNA after RNA transcript release at intrinsic terminators in a post-termination complex (PTC) *in vitro*. The RNAP in PTCs can diffuse in both directions on the DNA by sliding or hopping, and can flip 180 degrees on the DNA ^4–6^. The diffusing PTCs could sometimes initiate transcription in a promoter-dependent manner after reassociating with σ from solution, consistent with *in vivo* results ^4^. Available data indicate that complexes with the same properties as PTCs can be reconstituted by incubating core RNAP directly with DNA (reconstituted-PTCs, or rPTCs) ^4,7,8^.

The Sw2/Snf2 ATPase RapA is widespread throughout bacteria and is expressed to equal abundance as σ^70^ in *Eco* ^9,10^. RapA binds to core RNAP (Kd ∼5-10 nM) but not Eσ^70^ ^9,11^. Binding of RapA to RNAP strongly stimulates RapA ATPase activity, but DNA alone neither binds strongly to RapA nor stimulates its ATPase activity ^7,9–11^. While *rapA* null mutants do not display a clear *in vivo* phenotype in rich medium, stresses such as osmotic shock, exposure to sodium deoxycholate, or salt stress produce a protracted recovery phase in *rapA* knockouts ^12,13^. Additionally, *rapA* deletion diminishes antibiotic resistance in biofilms ^14^. Lastly, in *Vibrio cholerae*, deletion of the *rapA* homolog produces a >1000-fold decrease in colonization proficiency when cells attempt to engage the acid-tolerance response, which is important for virulence ^15^.

Many cellular functions have been proposed for RapA ^16–21^. In *in vitro* transcription reactions, RapA strongly stimulates multi-round transcript production, suggesting that it acts following termination ^12,16,22–24^. A recent single-molecule analysis defined the molecular target for RapA in the transcription cycle as the PTC; RapA forms a transient RapA-PTC complex that rapidly (within seconds) resolves upon ATP hydrolysis, releasing RapA and the RNAP from the DNA ^7^. Although several structural studies have provided insights into the interactions between RapA and core RNAP or the RNAP elongation complex (EC) ^18,20–22^, the structural mechanism for RapA function remains elusive in part because a structure of RapA with its true molecular target, the PTC, has not been determined.

Here, we structurally and biochemically characterize *Eco* rPTCs. We find that core RNAP can associate with negatively-supercoiled DNA, is capable of generating a transcription-competent transcription bubble *de novo*, and can initiate RNA synthesis at physiological nucleotide concentrations, all in the absence of a σ factor. We find that rPTC- mediated σ-independent transcription generates R-loops, potentially contributing to mutagenesis and posing a threat to genome stability *in vivo* ^25^. We show that RapA function *in vivo* plays a role in controlling R-loop formation. We also used a non-hydrolyzable ATP analog, ADP-AlF_3_ ^26^, to trap RapA associated with its functional target, the rPTC. A cryo-EM structure of RapA(ADP-AlF_3_)-rPTC reveals that RapA acts to impart a large conformational change in the RNAP that promotes release from the DNA. These findings have important implications for the canonical bacterial transcription cycle and the role played by PTCs and RapA in anti-sense and/or pervasive transcription and in genome stability ^27^.

## Core RNAP can associate with DNA and form a transcription bubble

For structural analysis of rPTCs, we sought to eliminate one-dimensional diffusion of the RNAP off the end of the DNA ^4^ as well as binding of RNAP to duplex DNA ends (end-binding) ^28^. We therefore assembled rPTCs by combining *Eco* core RNAP with a negatively-supercoiled circular DNA template, incubating briefly at 37°C, then prepared grids for cryo-EM analysis (Fig. 1a).

**Fig 1.**
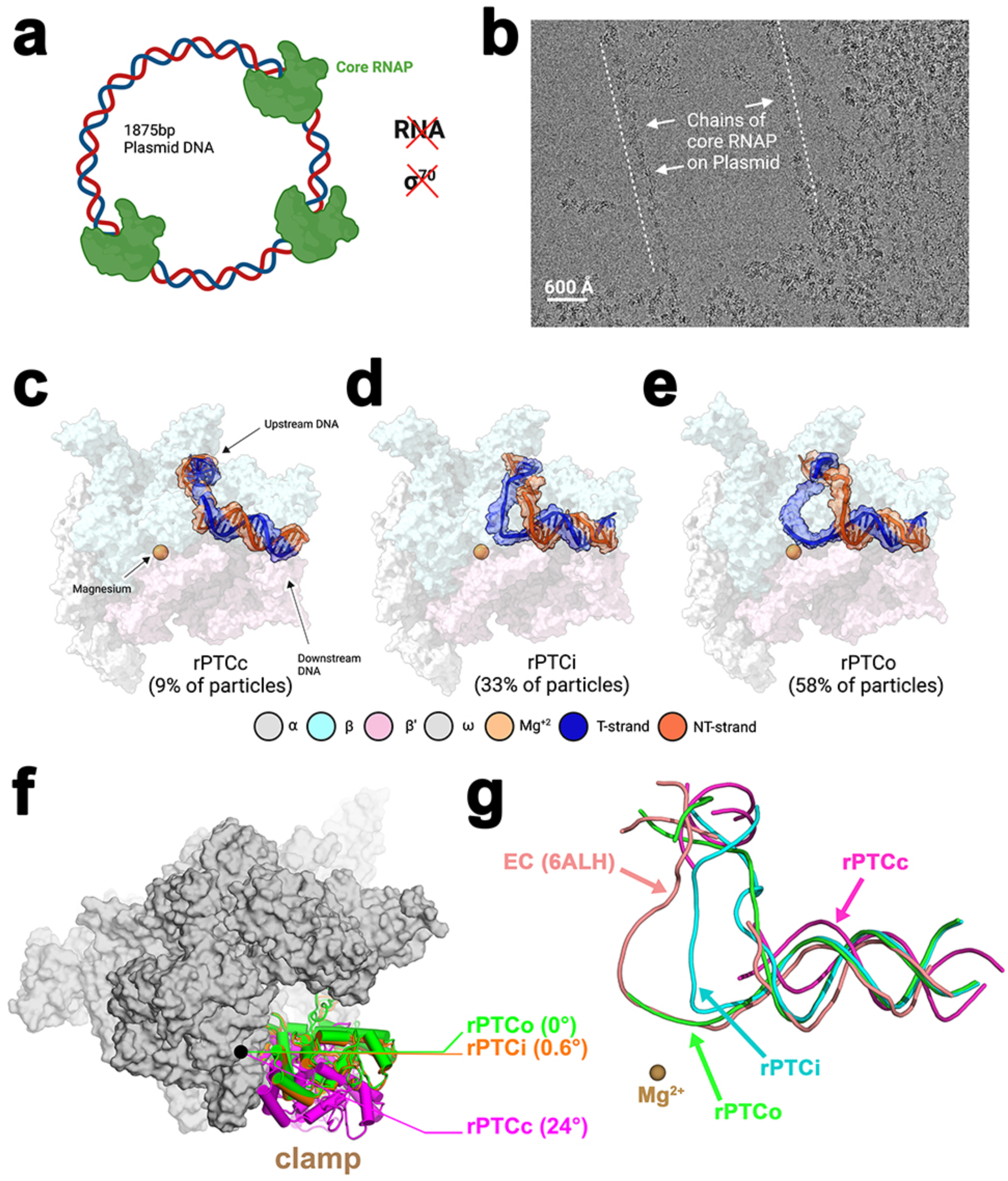
Reconstituted-PTCs unwind negatively-supercoiled DNA. **a.** Schematic illustrating the experimental setup. *Eco* core RNAP (green) was incubated with circular, negatively-supercoiled plasmid DNA in the absence of RNA, NTPs, and σ^70^. **b.** Representative micrograph illustrating core RNAP molecules associated with DNA. **c.-e.** Structures of rPTCs, determined by cryo-EM. The core RNAP is shown as a transparent molecular surface, revealing the DNA and active-site Mg^2+^ in the RNAP active-site cleft. The DNA is shown in cartoon format along with transparent cryo-EM difference density. **c.** rPTCc. **d.** rPTCi. **e.** rPTCo. **f.** RNAP clamp conformations for the rPTC structures. The rPTCo structure was used as a reference to superimpose rPTCc and rPTCi via α-carbon atoms of the RNAP structural core, revealing a common RNAP structure (shown as a grey molecular surface) but with clamp conformational changes characterized as rigid body rotations about a rotation axis perpendicular to the page (denoted by the black dot). The clamp modules are shown as backbone cartoons with cylindrical helices. The angles of clamp opening are shown relative to rPTCo (0°). **g.** Superposed DNAs from rPTCc (magenta), rPTCi (cyan), rPTCo (green), and an active EC (orange). The RNAP active site Mg^2+^ is shown as a brown sphere.

Visual inspection of micrographs revealed RNAP molecules associated with DNA, including segments of extended DNA decorated with chains of RNAP molecules like beads on a string (Fig. 1b). In our processing of the micrographs, we were unable to identify particles without DNA bound in the RNAP cleft (Extended Data Fig. 1). Steps of maximum likelihood classification ^29^ revealed three distinct conformational classes (Figs. 1c-f, Extended Data Figs. 1 and 2; Supplementary Data Table 1).

In the closed rPTC (rPTCc, ∼9% of the particle population; 4.7 Å nominal resolution; Fig. 1c, Extended Data Figs. 1 and 2a-c), the RNAP clamp ^30,31^ was notably wide open, 24° open compared to a 0° reference (Fig. 1f). The RNAP cleft was occupied with severely kinked (∼90°) but mostly duplex (closed) DNA (Fig. 1c). The vicinity of the DNA kink was poorly resolved and potentially a site of bubble nucleation, with the conserved Switch 2 (Sw2) and Fork Loop 2 (FL2) RNAP structural elements ^31^ proximal to the kink (Extended Data Fig. 3a). The RNAP β’rudder was well-resolved, situated in the major groove of the upstream double-stranded DNA (dsDNA) (Extended Data Fig. 3a).

In the intermediate rPTC (rPTCi; 33% of the particle population; 3.8 Å nominal resolution; Fig. 1d, Extended Data Figs. 1 and 2d-f), the RNAP clamp was relatively closed (0.6°; Fig. 1f). The RNAP cleft was occupied by DNA with a clear melted bubble of ∼5 nucleotides (nts) enclosed completely within the cleft (Fig. 1d). The single-stranded non-template strand (nt-strand) DNA was relatively well-resolved, but the template-strand (t-strand) was dynamic, poorly-resolved, and not near active site (Fig. 1g, Extended Data Fig. 3b).

The downstream fork junction of the rPTCi bubble appears to be stabilized by the insertion of Sw2 and FL2 elements between the melted strands at the downstream edge of the bubble (Extended Data Fig. 3b). The stabilized downstream fork junction observed together with the Sw2-FL2 insertion is consistent with prior biochemical findings implicating Sw2 in DNA melting, initiating-nucleotide binding, and promoter escape, along with EC stability ^32,33^. The β’rudder, previously implicated in EC and RPo stability ^34^, was disordered, owing to clash between the rudder and the duplex DNA immediately upstream of the growing bubble (Extended Data Fig. 3b).

In the open rPTC (rPTCo, 58% of the particle population; 3.6 Å nominal resolution; Fig. 1e, Extended Data Figs. 1 and 2g-i), the RNAP clamp was closed on the DNA (Fig. 1f). The RNAP cleft was occupied by DNA with a more extensive bubble of ∼7-8 nt (the upstream edge of the bubble was poorly-resolved), which propagated in the upstream direction compared with the smaller bubble in rPTCi (Figs. 1d and 1e). As in rPTCi, the single-stranded nt-strand DNA was relatively well-resolved while the t-strand was dynamic and poorly-resolved but occupied positions near the RNAP active-site Mg^2+^, similar to a *bona fide* elongation complex (EC; Fig. 1g).

As in rPTCi, the Sw2 and FL2 elements of rPTCo were inserted between the melted stands at the downstream fork of the transcription bubble (Extended Data Fig. 3c). The β’rudder was resolved and fully inserted between the two melted strands, likely stabilizing the upstream edge of the bubble (Extended Data Fig. 3c).

Rotation of a swivel module is associated with paused elongation complexes and appears to inhibit RNAP motions required to complete NTP binding, catalysis, and translocation ^35–38^.

Compared to an unswiveled reference structure (8EG8) ^38^, both rPTCi and rPTCo were significantly swiveled (4.4° and 4.0°, respectively; Extended Data Fig. 4).

## Core RNAP can initiate transcription on negatively-supercoiled DNA

Given the DNA disposition in rPTCo and the possibility that the t-strand DNA can occupy a position near the RNAP active-site similar to an active EC (Fig. 1g), we compared the transcription activity of the rPTCs with that of Eσ^70^ on a circular DNA plasmid template containing the strong T7A1 promoter and T7 intrinsic terminator sequence, expected to yield a specific 161 nt transcript with Eσ^70^ (Fig. 2a). We used relatively high NTP concentrations (500 μM each) to approximate physiological concentrations ^39^. The resulting RNA transcripts were extracted, quantitated, and the size distributions analyzed.

**Fig 2.**
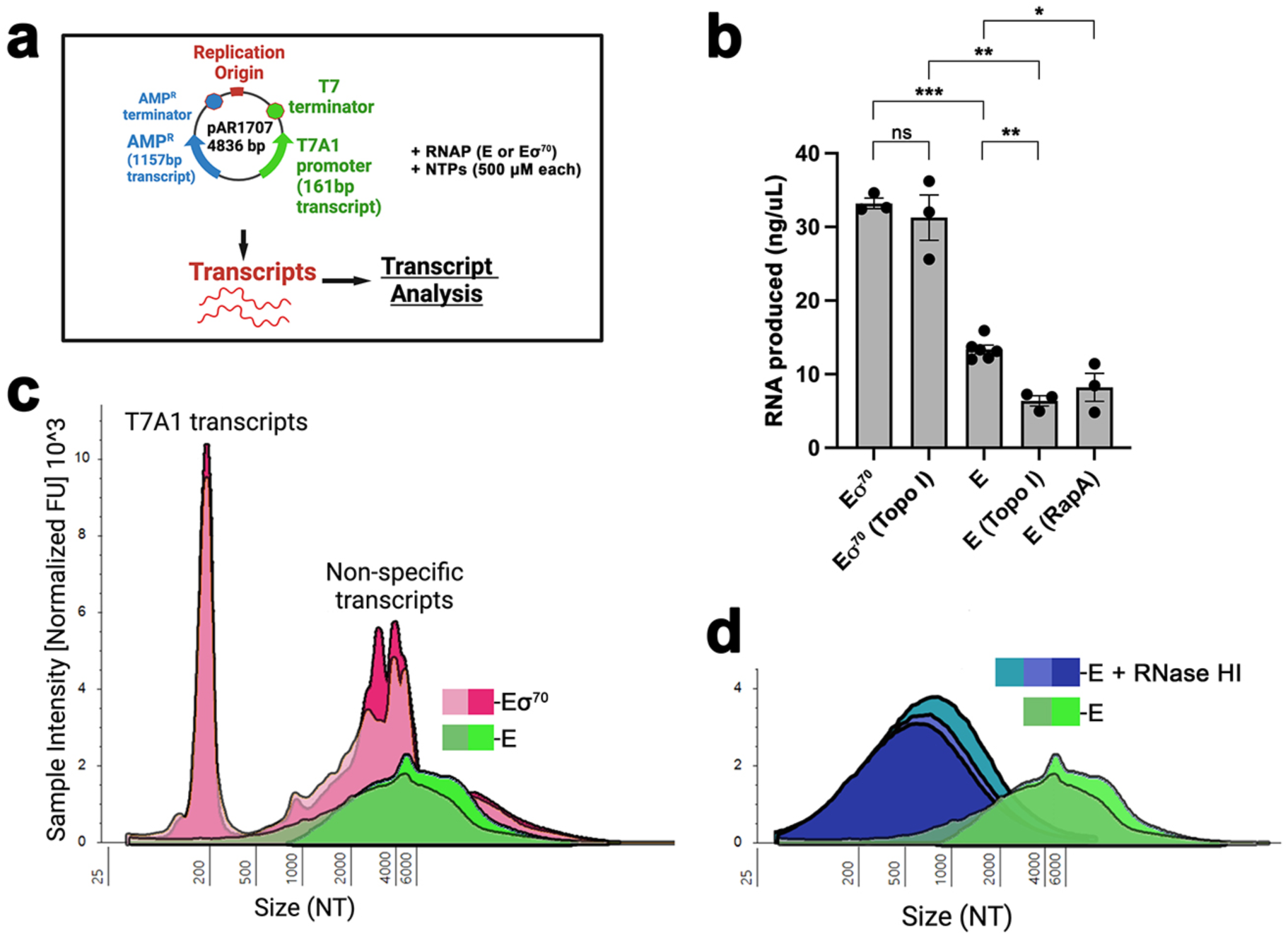
Transcription initiation by core RNAP. **a.** Schematic illustrating the experimental setup. *Eco* core RNAP (E) or Eσ^70^ was incubated with circular, negatively-supercoiled plasmid DNA (paAR1707)^69^ in the presence of NTPs (500 μM each) for 15 min at 37°C. The total amount and size distribution of the resulting RNA transcripts were then analyzed. **b.** Histogram plot showing the total amount of RNA produced from each transcription reaction. The bars denote the average of three to six independent measurements (individual data points shown). Error bars denote standard error. Statistical significance of differences between samples was determined using unpaired, two-tailed *t*-test (ns, *P* > 0.05; *, *P* ≤ 0.05; **, *P* ≤ 0.01; ***, *P* ≤ 0.001). **c.** Size distribution of transcripts resulting from two independent Eσ^70^ transcription reactions (red shades) and two independent core RNAP reactions (green shades). The area under the curve for each profile was normalized according to the total RNA produced. **d.** Size distribution of transcripts resulting from the core RNAP reactions (green shades) along with similar reactions treated with RNase HI (blue shades). The area under the curve for each profile was normalized according to the total RNA produced.

The overall amount and size distribution of the RNA produced by core RNAP vs. Eσ^70^ were clearly distinct (Figs. 2b and 2c), confirming the absence of significant σ^70^ contamination in our purified core RNAP. As expected, the transcription reactions with core RNAP produced less total RNA than Eσ^70^ (about 3-fold less; Fig. 2b). Reactions using plasmid DNA pre- incubated with topoisomerase I to relax supercoils showed a marked (more than 2-fold) decrease in core RNAP transcription whereas Eσ^70^ transcription was unaffected (core RNAP transcription on the topoisomerase I-treated template was ∼6-fold less than Eσ^70^; Fig. 2b). Transcription from the T7A1 promoter has been shown to be relatively insensitive to DNA supercoiling ^40^. The decreased core RNAP transcription on relaxed DNA could be explained by a decrease in the overall number of PTCs, or a decrease in the transcription activity of the PTCs. Since PTCs on relaxed DNA are extremely stable (Kd < 15 nM) ^8^ we favor the hypothesis that negative supercoiling shifts the equilibrium away from transcriptionally inactive PTCc and PTCi and towards transcriptionally active PTCo (Figs. 1c-e).

The size profile of the RNA products for the Eσ^70^ reactions was dominated by the expected specific transcript of 161 nt (Fig. 2c). Additional smaller peaks were superimposed onto a broad distribution of RNAs spanning approximately 1 kb to > 6 kb in length (Fig. 2c). By contrast, the core RNAP transcription products were characterized by a broad, relatively featureless distribution of RNAs over similar RNA lengths (1 kb to > 6 kb; Fig. 2c), consistent with non-specific transcription initiation by the rPTCs. These results clearly indicate that rPTCs can non-specifically initiate transcription, consistent with the cryo-EM structure of rPTCo that shows a complex containing a bubble with the t-strand loaded into the active site (Fig. 1e).

## PTC-initiated transcription is prone to R-loop formation

We noted that the broad distribution of large RNAs (> ∼1 kb) for both Eσ^70^ and core transcription reactions contained significant amounts of RNA chains longer than the plasmid itself (4.836 kb). We hypothesize that non-specific rPTC initiation generates elongating complexes with R-loops (persistent RNA/DNA hybrid) in their wake. The upstream RNA/DNA hybrid would prevent the formation of RNA secondary structure in the upstream transcript (such as terminator hairpins), causing the RNAP to ignore terminators and continuously transcribe around the circular DNA template in a rolling-circle transcription mechanism (*Eco* RNAP can produce > 7 kb transcripts from circular DNA templates in this manner) ^41^.

Treatment of the PTC transcription reactions with *Eco* RNase HI, which specifically hydrolyzes the RNA phosphate backbone when the RNA is hybridized to DNA, significantly reduced the size distribution of the RNAs (Fig. 2d), confirming extensive production of R-loops. The RNase HI treatment reduced the peak of the RNA size distributions by about 4.8 ± 0.1 kb, the size of the circular DNA template (Fig. 1a). This result supports the hypothesis that the core RNAPs transcribed all the way around the plasmid, generating an RNA/DNA hybrid the length of the DNA template. Further transcription would displace the RNA from the DNA template, allowing the production of RNA transcripts longer than the DNA template but always leaving an RNA/DNA hybrid the length of the DNA template.

## RapA contributes to the control of cytotoxic R-loops *in vivo*

Excessive PTCs on the DNA can lead to non-specific transcription initiation and generate extensive R-loops (Fig. 2d). Excessive R-loop formation *in vivo* is a threat to genomic stability and can be lethal ^25,42–44^. We hypothesize that RapA may contribute to the control of R-loops *in vivo* by removing PTCs, a potential source of R-loops. We first tested this hypothesis by comparing the growth of an *Eco rapA* null mutant (Δ*rapA*) with the wild-type parent strain (wt) under conditions of R-loop stress (Extended Data Fig. 5), induced by growing the cells in the presence of bicyclomycin (BCM), a selective inhibitor of the Rho termination factor ^45^. Rho plays an essential role in *Eco* by suppressing R-loop formation ^43,46^. The cells carried pBAD18*rnhA* (a plasmid expressing *rnhA*, encoding *Eco* RNase HI, under control of the arabinose-inducible P_BAD_ promoter) ^47^ or the empty pBAD18 plasmid as a negative control. We reasoned that a growth defect of the Δ*rapA* strain under R-loop stress would be compensated by overexpression of RNase HI via induction of pBAD18*rnhA*, directly confirming the role of R-loops. We analyzed cell growth (monitored by OD_600 nm_) using two parameters, the doubling time during log-phase growth and t_1/2_, the time for the cells to reach half their OD_600 nm_ plateau (Extended Data Figs. 5a-c).

In the absence of BCM, the Δ*rapA* strain had a small but reproducible growth defect (less than 1.2-fold) compared with wt under all conditions tested (Extended Data Figs. 5d and 5e). The growth rate defect of Δ*rapA* (relative to wt) increased significantly (doubling time, 1.3-fold; t_1/2_, 1.7-fold) when the cells were grown in the presence of BCM at 0.5X MIC (Figs. 3a and 3b, Extended Data Figs. 5f and 5g). The increased growth defect of Δ*rapA* in the presence of BCM was corrected by the expression of RNase HI (Figs. 3a and 3b, Extended Data Figs. 5f and 5g).

**Fig 3.**
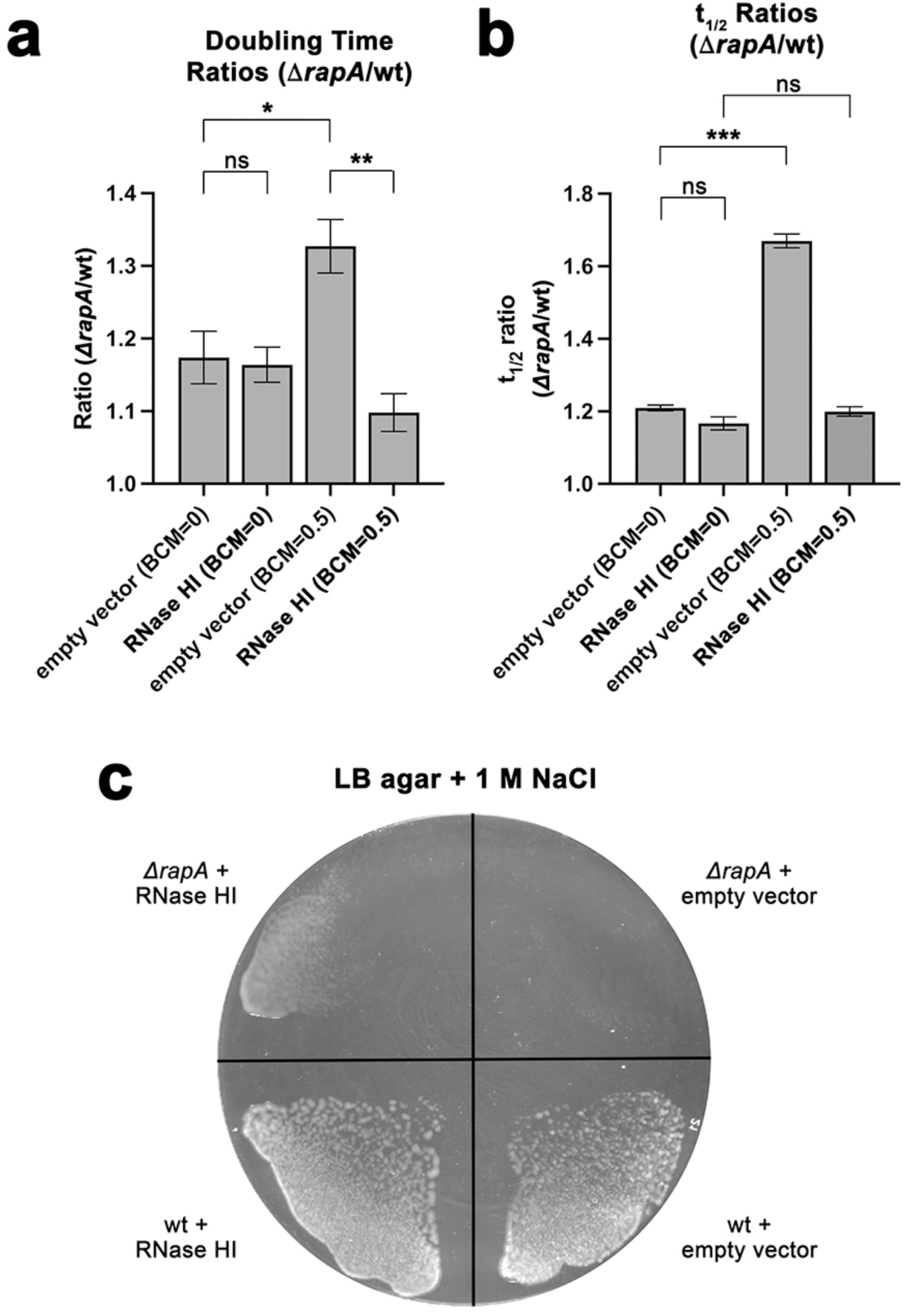
RapA suppresses cytotoxic R-loops *in vivo*. **a.-b.** Growth parameters (doubling times and t_1/2_) for wt and *ΔrapA Eco* cells carrying pBAD18 (empty vector) ^70^ or pBAD18*rnhA* (RNase HI) ^47^ without (BCM=0) or with 0.5X MIC BCM (BCM=0.5) were determined (Extended Data Fig. 5). Histograms show the ratios (*ΔrapA*/wt). Error bars denote standard error (≥ N=3 measurements). Statistical significance of differences between samples was determined using an unpaired, two-tailed *t*-test (ns, *P* > 0.05; *, *P* ≤ 0.05; **, *P* ≤ 0.01; ***, *P* ≤ 0.001). **a.** Ratios of doubling times at matching conditions (Δ*rapA*/wt). **b.** Ratios of t_1/2_ at matching conditions (Δ*rapA*/wt). **c.** The *ΔrapA* mutant cannot grow on LB plates + 1 M NaCl (upper right quadrant), but overexpression of RNase HI enables growth (upper left quadrant).

To test the hypothesis that RapA contributes to the control of R-loops *in vivo* at a distinct growth condition and without the use of BCM, we took advantage of a previous finding that deletion of *rapA* renders *Eco* unable to grow on LB plates with 1 M NaCl (Fig. 3c) ^12^. The apparent lethality of 1 M NaCl to Δ*rapA* was rescued by expression of RNase HI (Fig. 3c). This points to a role for R-loop toxicity in the inability of Δ*rapA* to grow on 1 M NaCl.

*Eco* responds to osmotic stress (such as 1 M NaCl in the surrounding medium) by accumulating high concentrations of osmolytes in the cytoplasm; major osmolytes include K^+^- ion, glutamate, and trehalose ^48,49^. These high concentrations of osmolytes can have significant effects on protein-DNA interactions ^50^. We suggest that the altered cytoplasmic conditions under osmotic stress may stabilize PTCs but that PTC accumulation is prevented by RapA activity in wt *Eco*. In the absence of RapA, PTCs accumulate, leading to increased levels of PTC-mediated initiation and production of R-loops to a lethal level.

## Cryo-EM structure of RapA(ADP-AlF_3_)-rPTC

Recent work identified PTCs as the target of the Swi2/Snf2 ATPase RNAP-recycling factor RapA ^7^. Inlow et al. ^7^ used single-molecule analyses to identify two kinetically distinct RapA- rPTC assemblies formed during ATP-dependent RNAP recycling (denoted RapA-PTC and RapA^†^-PTC) and proposed a kinetic scheme in which these assemblies are intermediates in disruption of PTCs by RapA (Extended Data Fig. 6a). Initial RapA association with PTCs was independent of nucleotide in solution, consistent with the first intermediate, RapA-PTC, containing apo-RapA (Extended Data Fig. 6a). Cryo-EM structures of apo-RapA engaged with core RNAP (7MKQ ^18^) and with ECs (7MKN ^18^, 7M8E ^21^) all yielded closed-clamp RNAP structures bound to apo-RapA in a nearly identical pose with each other, suggesting that the configuration of the complex of apo-RapA with RNAP is independent of the nucleic acid binding status of the RNAP. We therefore propose that the apo-RapA-RNAP and apo-RapA-EC structures represent good models for the structure of the first RapA-PTC intermediate.

The second kinetically distinct intermediate, RapA^†^-PTC, leads to PTC disruption at a rate that is strongly dependent on nucleotide in solution ^7^. In the presence of ATP, PTC disruption by RapA was at least 150-fold faster than when ATP was absent or when a non-hydrolyzable ATP analog was present. In the presence of the analog, RapA^†^-PTC was >20-fold more stable than the first intermediate, and more than half of the assemblies exhibited a characteristic lifetime close to 5 min ^7^, a time scale compatible with cryo-EM grid preparation. Therefore, we generated rPTCs on negatively-supercoiled circular DNA (Fig. 1a) and subsequently introduced RapA complexed with the non-hydrolyzable ATP analog ADP-AlF_3_ ^26^ and analyzed the resulting complexes by single-particle cryo-EM (Extended Data Fig. 6b). The results revealed a RapA(ADP-AlF_3_)-PTC structure (3.6 Å nominal resolution; Fig. 4a, Extended Data Fig. 6b and 6c; Supplementary Data Table 1) with novel characteristics compared to previously described RapA-RNAP or RapA-EC complexes ^18,21^ and that we equate with RapA^†^- PTC (Extended Data Fig. 6a). These novel properties inlude: i) the conformation of RapA, ii) the overall disposition of RapA with respect to the RNAP, and iii) the conformation of RNAP and the associated DNA. Cryo-EM density in the RNAP cleft of RapA^†^-PTC was consistent with kinked duplex DNA (Fig. 4a), very similar to the DNA occupying the rPTCc RNAP cleft (Fig. 1c).

**Fig 4.**
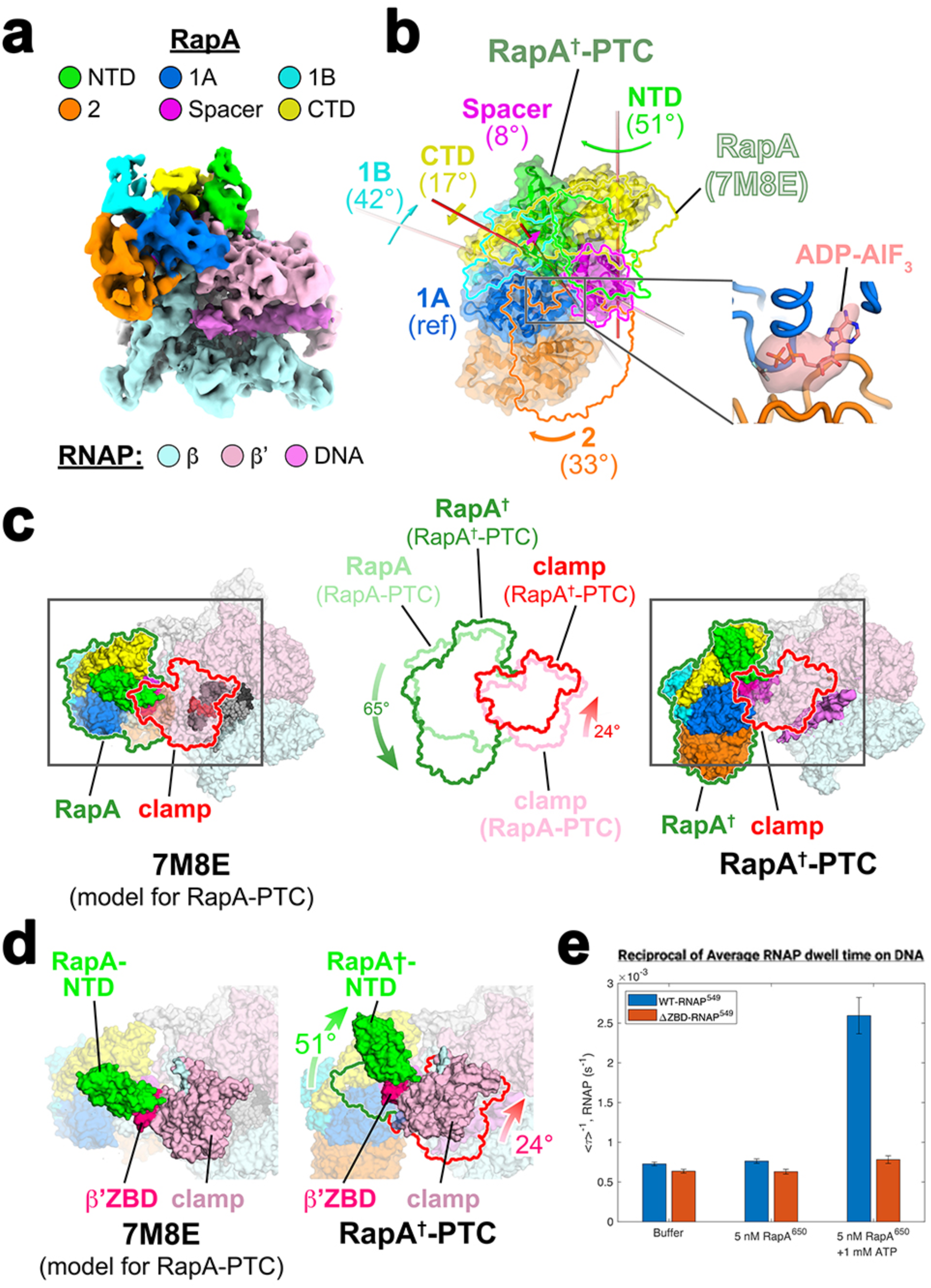
RapA opens the RNAP clamp to promote DNA dissociation. **a.** Cryo-EM map (local-resolution filtered) ^71^ of RapA(ADP-AlF_3_)-rPTC (RapA^†^-PTC). **b.** RapA from apo-RapA-PTC (using 7M8E as a model) ^21^ and RapA^†^-PTC were superimposed via the 1A domain (Supplementary Data Table 2). The RapA^†^ structure is shown in cartoon format with transparent molecular surfaces; the RapA domains are shown in colored outlines. The rotation and rotation axis for the conformational change of each domain (apo-RapA -> RapA^†^) is denoted. Note the large conformational change (51° rotation) of the RapA-NTD. The inset on the right shows a zoomed-in view of the nucleotide binding site (between the 1A and 2 domains). The modeled ADP-AlF_3_ and cryo-EM difference density corresponding to the bound nucleotide (transparent surface) is shown. **c.** Overall change in architecture of apo-RapA-PTC and RapA^†^-PTC. (*left*) apo-RapA-PTC is shown as a molecular surface (surfaces of RNAP are transparent to reveal the DNA inside the RNAP cleft). The downstream duplex DNA is shown as atomic spheres (t-strand, dark gray; nt-strand, light gray). RapA and the RNAP clamp are outlined in green and red, respectively. (*right*) RapA^†^-PTC is shown as a molecular surface (surfaces of RNAP are transparent to reveal the DNA inside the RNAP cleft). RapA and the RNAP clamp are outlined in green and red, respectively. (*middle*) The RapA-RNAP clamp outlines from the apo-RapA-PTC (lighter shades) and RapA^†^- PTC are superimposed. The RapA molecule as a whole rotates 65° on the surface of the RNAP, and the clamp opens 24°. **d.** Boxed regions of (**c**) are magnified, highlighting the RapA-NTD:β’ZBD interface. (*left*) apo-RapA-PTC (*right*) RapA^†^-PTC. The green and red outlines denote the positions of the RapA-NTD (green) and the RNAP clamp (red) from apo-RapA-PTC. A 51° rotation of the RapA^†^-NTD results in a 24° opening of the clamp. **C.** Results from single-molecule fluorescence microscopy experiments measuring the effective dissociation rates (reciprocal of the average RNAP dwell time on DNA, <τ>^-1^) of surface- tethered rPTCs formed with RNAP or with a ΔZBD-RNAP mutant, by RapA plus ATP or in controls lacking RapA or lacking ATP. Number of complexes from left to right: N=308, N=184, N=306, N=122, N=272, N=130. The error bars denote standard error.

## RapA binding to ADP-AlF_3_ triggers global conformational changes in RapA

The cryo-EM density of RapA^†^-PTC clearly showed ADP-AlF_3_ occupying the RapA nucleotide binding site between the 1A and 2 (RecA) domains (Fig. 4b; the RapA domains discussed herein are defined in Supplementary Data Table 2). As expected for a RecA-type ATPase ^51^, occupancy of the nucleotide binding site gave rise to a large change in the orientation of the RecA ATPase domains with respect to each other (RapA domains 1A and 2, Fig.4b) compared to apo-RapA structures ^18,21^, corresponding to a 33° rotation of domain 2 towards domain 1A (Fig. 4b). This large conformational change induced by ADP-AlF_3_ binding triggered a complex series of allosteric conformational changes in the other RapA domains (with respect to the reference domain 1A; Fig. 4b, Supplementary Video 1), most notably resulting in a motion of the RapA-NTD corresponding to a 51° rotation and a 27 Å translation of the domain center of mass (Fig. 4b).

## RapA conformational changes mechanically open the RNAP clamp

Comparing apo-RapA-EC (7M8E ^21^) with RapA^†^-PTC, the binding of ADP-AlF_3_ induced significant conformational changes in RapA (Fig. 4b, Supplementary Video 1). These changes were accompanied by a large rearrangement of RapA as a whole, corresponding to a 65° rotation of RapA relative to the RNAP (Fig. 4c, Supplementary Video 2). Major RapA-RNAP interfaces include the RecA domains (RapA domains 1A and 2; Fig. 4b) with the RNAP βflap-tip and the RapA-NTD with the RNAP β’ZBD (Figs. 4c and 4d).

The main anchor point for the 65° RapA rotation (with respect to the RNAP) is the RapA-1A:βflap-tip interaction. The rotation of RapA about this anchor point, which is accommodated by flexibility of the βflap-tip (Extended Data Fig. 7a, Supplementary Video 2), results in the large motion (51° rotation, 27 Å translation) of the RapA-NTD mentioned previously (Figs. 4b and 4c). The RapA-NTD forms a significant interface with the β’ZBD [583 Å^2^ interface area ^52^]; consequently, the large motion of the RapA-NTD pulls the β’ZBD and associated RNAP clamp with it, resulting in a 24° opening of the RNAP clamp (Figs. 4c and 4d, Supplementary Video 2). The RapA-Spacer domain also appears to wedge itself into the now open RNAP cleft, possibly stabilizing the open-clamp conformation (Extended Data Fig. 7b).

The relatively closed clamps observed in the rPTCo and rPTCi structures were accompanied by DNA melting observed in those structures (Figs. 1d and 1e), whereas rPTCc, with its open clamp, contained apparently closed, duplex DNA in its RNAP cleft (Fig. 1c).

Similary, the cryo-EM density in the RNAP cleft of the open-clamp RapA^†^-PTC is consistent with duplex DNA (Fig. 4a). Clamp opening in RapA^†^-PTC would not only allow the DNA to anneal into a duplex but would also be expected to destabilize DNA binding and promote dissociation of the DNA. Consistent with this expectation, we observed that the addition of RapA (in the presence of ATP) to our *in vitro* core RNAP transcription reactions inhibited overall RNA synthesis by nearly 2-fold compared to core RNAP alone (Fig. 2b).

## RapA uses its NTD to pull on the β’ZBD to open the RNAP clamp

Our model for RapA function predicts that the RapA-NTD:β’ZBD interface is crucial, allowing the motion of the RapA-NTD to pull on the β’ZBD and open the RNAP clamp. We tested this model by assessing the ability of RapA to disrupt PTCs generated from an RNAP derivative lacking the β’ZBD (ΔZBD-RNAP). Non-supercoiled fluorescently-labeled DNA circles tethered to the surface of a glass flow chamber were preincubated with either 1.5 nM RNAP or 4 nM ΔZBD-RNAP, each dye-labeled via a SNAP-tag fusion to the β′ subunit C-terminus (RNAP^549^ and ΔZBD-RNAP^549^, respectively). Single-molecule total internal reflection microscopy (smTIRF) revealed individual molecules of RNAP at the locations of single DNA molecules, indicating rPTC formation. Note that a higher concentration of ΔZBD-RNAP^549^ was required to achieve quantities of rPTCs optimal for experiments; at 1.5 nM RNAP^549^, we attained 40-50% DNA occupancy by RNAP, compared to ∼25% occupancy with 4 nM ΔZBD-RNAP^549^.

As previously reported ^7^, replacement of the flow chamber solution with a buffer containing 1 mM ATP and a fluorescent RapA derivative (5 nM RapA^6^^50^) accelerated the loss of RNAP from the surface compared to controls without RapA or without ATP (Fig. 4e), indicating rPTC disruption. By contrast, no disruption above control was detected with ΔZBD-RNAP, highlighting the essential role of the interaction between the RapA-NTD and the RNAP-β’ZBD in stimulating RNAP clamp opening and subsequent dissociation of RNAP from DNA.

## Discussion

Both inter-genic and antisense transcription initiation are of growing interest as genome-wide sequencing techniques have allowed deeper coverage of *in vivo* transcription to be achieved ^53^. The role played by PTCs in the production of this promoter-independent transcription has been difficult to structurally probe due to challenging biochemical reconstitution constraints. PTCs exhibit one-dimensional diffusion on the DNA ^4–6^ (meaning they could slide off the ends of a linear DNA fragment) and core RNAP binds tightly to the ends of linear DNA fragments ^28^.

These features preclude the use of linear DNA fragments typically used in cryo-EM analyses of protein-DNA complexes. By reconstituting PTCs on a negatively-supercoiled circular duplex DNA substrate, our results provide insight into the structural nature of PTCs and promoter-independent transcription initiation more broadly, and the role of RapA in preventing the excessive build-up of PTCs, leading to a revised model for the bacterial transcription cycle (Fig. 5) ^4^.

**Fig 5.**
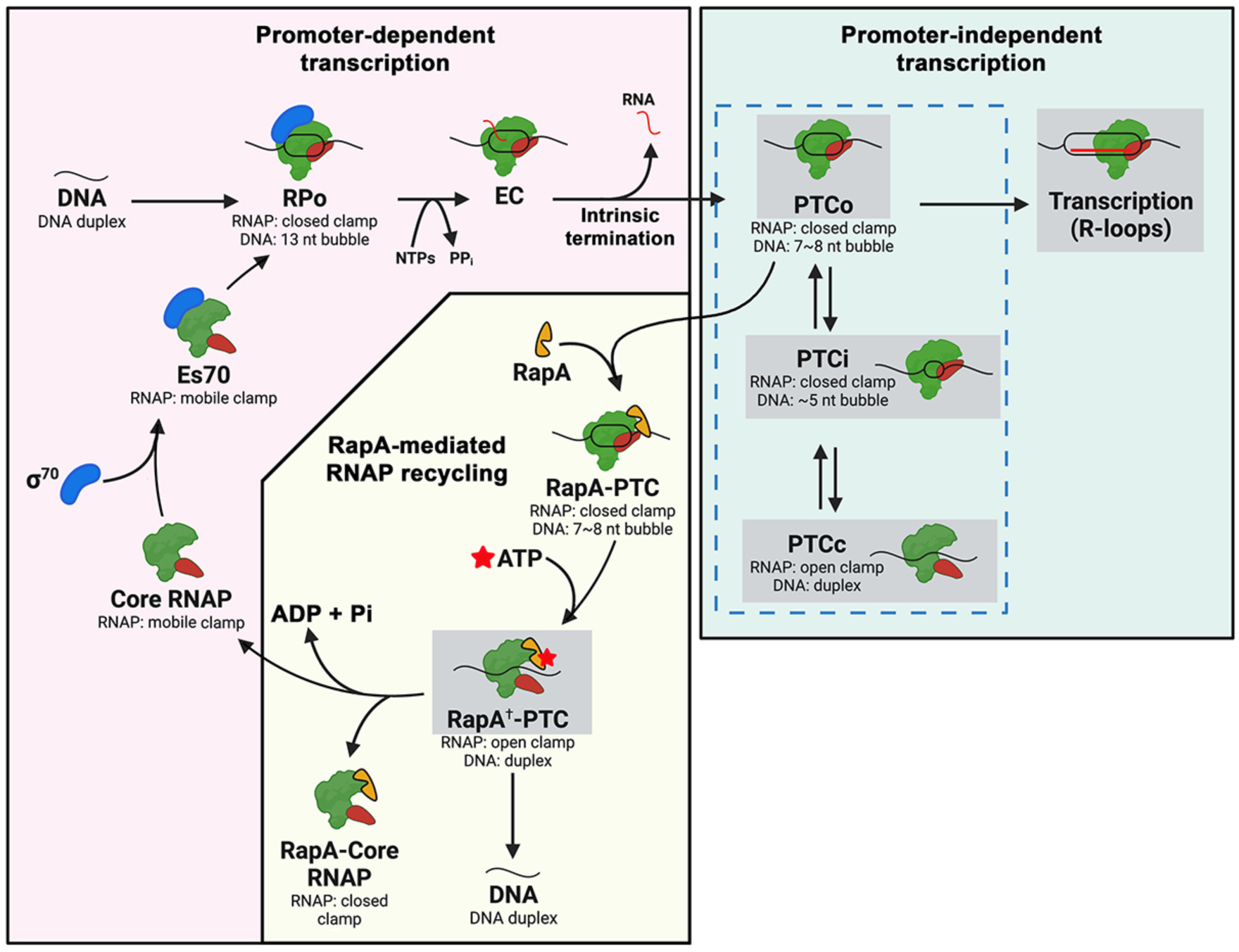
Revised model for the Bacterial Transcription Cycle. Results from this work (highlighted in gray boxes) yield a model of bacterial transcription where free core RNAP combines with σ^70^ to form Eσ^70^, which locates promoter DNA sequences and initiates promoter-specific transcription of RNA (promoter-dependent transcription, light pink background). Following intrinsic termination, core RNAP can remain on the DNA in a PTC, equilibrating between PTCo ⮀ PTCi ⮀ PTCc. PTCo can initiate transcription independent of σ but is prone to cytotoxic R-loop formation (promoter-independent transcription, light green background). This toxic component of the bacterial transcription cycle is suppressed by RapA, which specifically binds PTCs and pulls open the RNAP clamp in an ATP-dependent manner to allow the transcription-bubble to reanneal and remove the core RNAP from DNA, facilitating RNAP recycling and averting the harmful build-up of R-loops (RapA-mediated RNAP recycling, light yellow background).

σ-dependent transcription begins with base-flipping and capture of conserved nt-strand bases (normally at position -11 and -7 with respect to the transcription start site at +1) ^54^ within the -10 core promoter element, triggering transcription bubble nucleation and bubble propagation downstream to the transcription start site ^55,56^. Our rPTC structures show that core RNAP can bind and bend the DNA to nucleate a transcription bubble independent of σ factor or conserved promoter elements (Figs. 1c-f). Core RNAP appears to nucleate its transcription bubble from within the RNAP active-site cleft and to propagate in the upstream direction while engaging SW2, FL2, and the β’rudder in the reverse order from σ-dependent transcription initiation (Extended Data Fig. 3).

Evolved strategies to suppress promoter-independent transcription appear manifold. Termination factor Rho targets ECs producing aberrant transcripts ^57^. Additional mechanisms for preventing non-specific association of RNAP with DNA outside of promoter sequences include auto-inhibitory σ^70^_1.1_ ^58^, DNA packaging proteins (such as H-NS) ^59^, and RapA ^7^. Our approach for generating rPTCs suitable for cryo-EM analysis allowed us to visualize RapA engaged with its true target substrate (Fig. 4a). Our RapA^†^-PTC structure, along with comparisons to previously determined apo-RapA-RNAP and apo-RapA-EC structures, illustrate how nucleotide binding drives a complex allosteric conformational rearrangement of RapA that pulls open the RNAP clamp, allowing melted regions of the DNA to rewind and promoting dissociation of the DNA. Further work will be required to determine the role of ATP hydrolysis (as distinct from nucleotide binding) in the RapA functional cycle.

Dey et al. ^60^ found that a population of *Eco* RNAP ECs paused at the U-rich pause site following the HK022 putL element released their RNA transcript but remained associated with the DNA template - essentially PTCs. The RNAP conformation in these PTCs (8AC2) ^60^ closely matches the rPTCc RNAP conformation, and both of these are very similar to the PTC portion of our RapA^†^-PTC structure (Supplementary Data Table 3). Thus, RapA uses ATP binding energy to stabilize a pre-existing PTC state. Dey et al. ^60^ did not observe closed-clamp/open-bubble structures corresponding to rPTCi or rPTCo, presumably because their complexes were formed on linear DNA fragments that lacked supercoiling.

Our *in vitro* and *in vivo* results point to the formation of cytotoxic R-loops via σ^70^- independent transcription mediated by PTCs (Figs. 2d and 3). Our observation that σ^70^- independent transcription initiation is prone to R-loop formation points to a crucial role for σ^70^ in initiating proper RNA transcript strand separation from the RNA-DNA hybrid during initial transcription at promoters. The σ^70^-family σ factors ^1^ contain a conserved structural element, the σ-finger (also called the σ^70^_3.2_-loop) ^61,62^, that loops into the RNAP cleft and helps pre-organize the t-strand DNA near the active site but also blocks the path of the elongating nascent RNA ^61,62^. As the RNA chain extends during initiation, a steric clash with the σ-finger either promotes abortive initiation or the σ-finger is displaced, facilitating promoter escape ^61–64^. We hypothesize that the absence of the σ^70^-finger in the RNAP cleft results in unsuccessful strand separation of the RNA-DNA hybrid during PTC transcription initiation.

Our *in vivo* experiments show that overexpression of RNase HI rescues growth defects of a *rapA* null strain (Δ*rapA*) at two distinct growth conditions, R-loop stress induced by sub-MIC BCM (Figs. 3a and 3b, Extended Data Fig. 5) and osmotic stress induced with 1 M NaCl (Fig. 3c). As observed previously ^12^, the Δ*rapA* strain appeared to be completely unable to grow on LB agar + 1 M NaCl (Fig. 3c), suggesting that *rapA* may be essential under this condition (conditional essentiality). Strikingly, overexpressing RNase HI allowed growth at this otherwise lethal condition for the Δ*rapA* strain. This result strongly suggests that under osmotic stress, the loss of RapA function leads to a lethal accumulation of R-loops, presumably generated by uncontrolled PTC-mediated initiation.

Microbes are ubiquitous across a vast assortment of environments and consequently have developed survival strategies for frequent osmotic shock ^48^. *Eco* has effectively solved the biophysical challenge of surviving in aqueous environments ranging from highly dilute solutions to those containing molar concentrations of salts ^49^. Here, our results highlight the lethal threat posed by cytotoxic R-loops and how the activity of RapA in evicting PTCs permits cell survival.

All-in-all, our findings implicate transcription initiation by PTCs, either before or after the association of σ, in promoter-independent transcription and the generation of cytotoxic R-loops in bacteria. Additionally, our structure of the RapA^†^-rPTC delineates the mechanisms underlying the suppression of promoter-independent transcription by RapA. We reveal a previously unappreciated role for RapA *in vivo*; contributing to the suppression of deleterious R-loops. RapA is widely-spread among bacterial lineages, but not universally so. We hypothesize that bacterial lineages that lack RapA may harbor analagous ATPases that fulfill similar roles as *Eco* RapA ^65–68^. The behavior of bacterial RNAPs not related to *Eco* RNAP after intrinsic termination is unknown; further studies will be required to understand how the potential for promoter-independent transcription and R-loop accumulation is suppressed in bacteria that lack RapA.

## Methods

Structural biology software was accessed through the SBGrid consortium ^72^. No statistical methods were used to predetermine sample size. The experiments were not randomized. The investigators were not blinded to allocation during experiments and outcome assessment.

### Protein expression, purification, reconstitution, and and labeling

*Eco* core RNAP and σ^70^ were separately overexpressed and purified as previously described ^55^. *Eco* core RNAP fluorescently labeled with SNAP-Surface DY-549 dye (New England Biolabs) via a SNAP-tag on the C-terminus of β’ (WT-RNAP^549^) was expressed, purified, and labeled as described ^4^. A SNAP-tagged β’ zinc binding domain deletion mutant (ΔZBD-RNAP^549^; β’ residues 64-94 replaced with a -GS- linker) was similarly purified and labeled. Native mass spectrometry (nMS) analysis of unlabeled SNAP-RNAP samples showed that the WT-SNAP-RNAP was mostly assembled core (80%); by contrast, only 9% of β’-ΔZBD-SNAP-RNAP was fully assembled with 85% of the mutant core RNAP lacking the ω subunit. To avoid potential issues due to the low abundance of ω in the β’-ΔZBD-SNAP-RNAP sample, WT-SNAP-RNAP and ΔZBD-SNAP- RNAP were labeled with DY-549 dye as described ^4^ then incubated with three-fold and four-fold excess ω subunit (respectively) in the following steps: WT-RNAP and ΔZBD-RNAP were labelled with DY-549 dye as described ^4^. For ΔZBD-RNAP^549^, 1.83 μM ΔZBD-RNAP^549^ was mixed with 7.68 μM ω subunit and incubated on ice for 30 min. For WT-RNAP^549^, 12 μM WT- RNAP^549^ was mixed with 36 μM ω subunit and incubated on ice for 30 min. The samples were then buffer-exchanged into 10 mM Tris-HCl, pH 8, 100 mM NaCl, 2.5 mM MgCl_2_, 1 mM dithiothreitol (DTT), and 25% glycerol (v/v) using Princeton Separations Centri-Spin10 (5 kDa) spin columns and stored at -80°C. Subsequent nMS analysis revealed that both WT-RNAP^549^ and ΔZBD-RNAP^549^ core complexes were fully assembled and completely labelled with DY-549.

*Eco* RapA was overexpressed and purified as previously described ^7^. A fluorescently-labelled SNAP-RapA construct, RapA^6^^50^, was prepared as previously desribed ^7^

### Native MS analysis

The RNAP samples were buffer-exchanged into nMS solution (500 mM ammonium acetate, pH 7.5, 0.01% Tween-20) using Zeba microspin desalting columns (Thermo Scientific) with a 40-kDa MWCO ^73^. For nMS analysis, 2–3 µL of the buffer-exchanged sample was loaded into a gold-coated quartz emitter that was prepared in-house and then electrosprayed into an Exactive Plus EMR instrument (Thermo Fisher Scientific) with a static nanospray source ^74^. The nMS parameters used included: spray voltage, 1.2 kV; capillary temperature, 125 – 150 °C; in-source dissociation, 10 V; S-lens RF level, 200; resolving power, 8,750 or 17,500 at *m/z* of 200; AGC target, 1 x 10^6^; maximum injection time, 200 ms; number of microscans, 5; injection flatapole, 8 V; interflatapole, 4 V; bent flatapole, 4 V; high energy collision dissociation (HCD), 200 V; ultrahigh vacuum pressure, 5.5–6.5 × 10^−10^ mbar; total number of scans, at least 100. Mass calibration in positive EMR mode was performed using cesium iodide. The acquired MS spectra were visualized using Thermo Xcalibur Qual Browser (v. 4.2.47) and processed further using the deconvolution software UniDec version 4.2.0 ^75,76^ to obtain the deconvolved masses. The resulting measured masses for the unlabeled SNAP-RNAP assemblies observed included WT core: 412,045 Da, WT core–ω: 402,035 Da, α_2_b: 223,691 Da, b’-DZBD core: 408,544 Da and b’-DZBD core–ω: 398,427 Da. The measured masses for the DY549-labeled, SNAP-RNAP assemblies incubated with excess ω subunit were WT-RNAP^549^: 413,105 Da and b’-DZBD-RNAP^549^: 409,687 Da, which closely matched the mass of the corresponding RNAP core with one covalently attached DY-549 dye. The mass accuracies, calculated as the percent mass deviation between the measured and predicted masses, ranged from 0.03% - 0.07%.

### Transcription Assays

Eσ^70^ was reconstituted by incubating core RNAP (0.5 μM final) with σ^70^ (2.5 μM final) at 37°C for 15 min. Either core RNAP or Eσ^70^ (as indicated) were incubated with ATP, CTP, GTP, and UTP (500 μM each; TriLink Biotechnologies) and pAR1707 plasmid (35.8 nM final; Figure 1A) ^69^ at 37°C for 15 min in transcription buffer (100 mM Tris-HCl, pH 8.0, 500 mM KCl, 100 mM MgCl_2_, 1 mM EDTA, 10 mM DTT, and 50 μg/mL BSA). Next, two units of Turbo DNase I (Invitrogen) were added to the sample along with Turbo DNase I reaction buffer (Invitrogen, final concentration 1X) and incubated at 37°C for 15 min in order to fully digest all DNA substrate present in the reaction, effectively halting transcription. DNase I digestion was stopped by adding EDTA to 15 mM (final) and RNA was extracted with RNA Clean and Concentrator Kit (Zymo Research). Eluted RNA was combined with Qubit HS RNA Reagent and Qubit HS RNA Buffer and measured using a Qubit fluorometer using the HS RNA protocol after RNA standardization (Life Technologies). The size profile of the RNA sample was analyzed on an Agilent 2200 TapeStation using a High sensitivity RNA Screentape with RNA sample buffer (Agilent). The results were visualized using the Agilent Tapestation Software (Agilent). For RNase H sensitivity experiments, transcription was halted prior to the introduction of Turbo DNase I by the addition and 5 min incubation of 5 μM Rifampicin at 37°C. Subsequently, 2.5 units of RNase H (New England Biolabs) was added to the reaction and incubated at 37°C for 5 minutes.

### Cloning of cryo-EM plasmid scaffold

To construct a high copy-number plasmid (pJB1) small enough to permit cryo-EM data collection, we performed Gibson Assembly on amplicons generated from plasmids pAR1707 and pUC57. The pAR1707 insert amplicon contains a T7A1 promoter sequence and downstream 21-mer stall sequence derived from pAR1707 (for use outside of the scope of this work). The pUC57 backbone amplicon included an Ampicillin resistance gene and a high copy-number origin of replication.

### Preparation of *Eco* rPTC complex for cryo-EM

*Eco* core RNAP (0.5 mL of 5 mg/mL protein) was injected into a 10/300 Superose 6 Increase column (Cytiva) equilibrated with 10 mM Tris-HCl, pH 8.0, 100 mM KCl, 5 mM MgCl_2_ and 2.5 mM DTT. The peak fractions of the eluted protein were concentrated by centrifugal filtration (EMD-Millipore - 30 kDa MWCO) to 25 μM protein concentration.

Plasmid pJB1 was grown overnight in DH5α cells in standard Luria broth with 100 μg/ml ampicillin and isolated using a Plasmid DNA Maxiprep kit (Qiagen). Plasmid DNA solution at 1750 ng/μL was added to core RNAP for a final concentration of 491 ng/μL of DNA (0.82 μM). The sample was incubated for 15 min at 37°C, then 3-([3- cholamidopropyl]dimethylammonio)-2-hydroxy-1-propanesulfonate (CHAPSO) was added to a final concentration of 8 mM ^77^ and the sample was kept at room temperature prior to grid preparation.

### Preparation of RapA-rPTC complex for cryo-EM

The rPTC complexes were prepared as described above but with the plasmid final concentration of 0.42 μM. RapA (20 μM) was pre-incubated with AlF_3_ and ADP (2.5 mM each) (Sigma-Aldrich). The ADP-AlF_3_-RapA solution was added to the rPTC sample to achieve a final concentration of 8 μM RapA, 1 mM AlF_3_, and 1 mM ADP. CHAPSO was then added (8 mM final concentration) and the sample was kept at room temperature prior to grid preparation.

### Cryo-EM grid preparation

C-flat holey carbon grids (CF-1.2/1.3-4Au; Protochips) were glow-discharged for 20 s before the application of 3.5 μL of the sample (0.42 μM Plasmid DNA, 8 μM core RNAP, 8mM Chapso, 8 μM RapA, 1 mM ADP, 1 mM AlF3, 8 mM CHAPSO). After blotting for 3–4.5 s the grids were plunge-frozen in liquid ethane using an FEI Vitrobot Mark IV (FEI) with 100% chamber humidity at 37°C.

### Cryo-EM data acquisition and processing

Eco *rPTCs*. Grids were imaged using a 300-keV Titan Krios (FEI) equipped with a K3 Summit direct electron detector (Gatan). Images were recorded with Leginon ^78^ in counting mode with a pixel size of 1.076 Å and a defocus range of -0.25 to -4.16 μm. Data were collected with a dose rate of 28 *e*^-^ per Å^2^ per s. Images were recorded over a 2 s exposure with 0.05 s frames (40 total frames) to give a total dose of 55.9 *e*^-^/Å^2^. Dose-fractionated videos were gain-normalized, drift-corrected, summed, and dose-weighted using MotionCor2 ^79^. The contrast transfer function (CTF) was estimated for each summed image using the Patch CTF module in cryoSPARC3 (CS3) ^80^. Particles were picked and extracted from the dose-weighted images with a box size of 256 px using CS3 Blob Picker and Particle Extraction. Coordinates pointing to contaminating ice particles were extracted as faux particles and used to generate an initial decoy 3D model in CS3 (ab initio reconstruction) in order remove junk particles from initial particle stacks. Multiple rounds of CS3 Hetero Refinement of all blob-picked particles employing 3D templates from this 3D decoy along with an *Eco* core RNAP 3D template (PDB: 6ALH with all nucleic acids removed, low pass filtered to 20 Å resolution), were used to identify a 3D consensus reconstruction containing subclasses for rPTCo and rPTCi. Multiple rounds of CS3 Hetero Refinement of all blob-picked particles employing 3D templates of the 3d decoy along with *E.* coli core RNAP 3D template (PDB: 6GH6 with all nucleic acids removed, low pass filtered to 20 Å resolution), were used to identify a 3D consensus reconstruction containing the subclass for rPTCc. Many classification schemes were tested that converged on the conclusion that three mid-to-high-resolution classes were present in the particle dataset. All three classes were subjected to two rounds of successive Bayesian Polishing in Relion3 ^81^. CS3 CTF-refinement and non-uniform (NU) refinement were then performed for each resulting class, yielding three distinct structures: rPTC_c_ (13,101 p, 4.7 Å nominal resolution), rPTC_i_ (49,701 p, 3.8 Å nominal resolution), and rPTC_o_ (86,865 p, 3.6 Å nominal resolution) (Figure S1).

*Eco rPTC + RapA.* Grids were initially screened using a 200 keV Talos Arctica (FEI) equipped with a K2 Summit direct electron detector. Datasets were recorded with a pixel size of 1.5 Å over a defocus range of −1.0 μm to −3.5 μm. Movies were recorded in counting mode at 8 electrons/physical pixel/second in dose-fractionation mode with subframes of 0.3 s over a 15 s exposure (50 frames) to give a total dose of 53.33 electrons/Å^2^. Dose-fractionated movies were gain-normalized, drift-corrected, summed, and dose-weighted using MotionCor2 ^79^. The CTF was estimated for each summed image using the Patch CTF module in CS3 ^80^. Particles were picked and extracted from the dose-weighted images with a box size of 256 px using CS3 Blob Picker and Particle Extraction. Particles were curated via CS3 2D classification and selection. CS3 ab initio reconstruction was used to produce a density map of rPTC + RapA. CS3 Non- uniform refinement was used to further refine this initial map to 6.16Å. Core RNAP subunits (no nucleic acids present) and all RapA domains, except for spacer domain, were rigid-body refined to fit the density of the map. The resulting incomplete molecular model was used to produce a simulated 20 Å resolution density map for downstream templating.

For our full data collection, grids were imaged using a 300-keV Titan Krios (FEI) equipped with a K3 Summit direct electron detector (Gatan). Images were recorded with Leginon ^78^ in counting mode with a pixel size of 1.076 Å and a defocus range of -0.8 to -2.5 μm. Data were collected with a dose rate of 26 *e*^-^ per Å^2^ per s. Images were recorded over a 2-s exposure with 0.05-s frames (40 total frames) to give a total dose of 56 electrons per Å^2^. Dose- fractionated videos were gain-normalized, drift-corrected, summed, and dose-weighted using MotionCor2 ^79^. The CTF was estimated for each summed image using the Patch CTF module in CS4 ^80^. Particles were picked and extracted from the dose-weighted images with a box size of 256 px using CS3 Blob Picker and Particle Extraction. An initial decoy 3D model was generated in CS4 (ab initio reconstruction) as described above. Multiple rounds of CS4 Hetero Refinement of all blob-picked particles employing 3D templates of this 3D decoy along with the incomplete 20 Å ab initio map mentioned above (all nucleic acids and RapA spacer domain missing), were used to identify a 3D consensus reconstruction containing rPTC+RapA. This class was subjected to focused CS4 3D-classification, masking around RapA to identify a clear class for RapA^†^-PTC. Then, two rounds of successive Bayesian Polishing were performed in Relion3 ^81^. Then, CS4 CTF-refinement and NU-refinement were performed, yielding RapA^†^-PTC from 100,010 p (3.6 Å nominal resolution) (Figure S5B).

The heatmap distributions of particle orientations and half-map FSCs were calculated using CS3. 3D Fourier shell correlation calculations were performed using 3DFSC ^82^. Local- resolution calculations were performed using blocres and maps were locally filtered using blocfilt (Bsoft package) ^71^.

### Model building and refinement

*rPTCs.* The initial model for the rPTCs was derived from PDB 8EG7 ^38^ with all of the nucleic acids removed. The model was manually fit into the cryo-EM density maps using ChimeraX ^83^ and rigid-body and real-space refined using PHENIX real-space-refine ^84,85^. For real-space refinement, rigid-body refinement was followed by all-atom and *B* factor refinement with Ramachandran and secondary structure restraints. Models were inspected and modified using COOT ^86^.

*RapA^†^-PTC.* The initial model for RapA^†^-PTC included the rPTCc model (determined herein) combined with RapA from PDB 7M8E ^21^. Steps of model building and refinement followed the same steps for the rPTCs described above.

### Single-molecule experiments

Construction of the 586 bp circular promoter-less DNA templates (npDNA^Cy5^) as well as the expression, purification, cloning, and labelling of RapA^6^^50^ were done as previously described ^7^. The workflow for the single-molecule fluorescence washout experiments and data analysis was performed as previously described ^7^ except that we introduced either 1.5 nM RNAP^549^ or 4 nM ΔZBD-RNAP^549^ into the chamber in transcription buffer, incubated for 10 min to allow for rPTC formation, and washed out excess unbound RNAP^549^.

Then, solutions of wash buffer, 5 nM RapA^6^^50^, or 5 nM RapA^6^^50^ + 1 mM ATP were mixed in transcription buffer and loaded into the glass chamber at time zero. Image acquisition began within 10 s after loading the reagents (at *t* = 0), with excitation alternating between 532 nm and 633 nm (400 µW each) at a frame rate of 1 frame/s for 40 min. Characteristic lifetimes of RNAP^549^ and ΔZBD-RNAP^549^ on npDNA^Cy5^ were calculated by taking the average <τ> of all measured intervals where RNAP^549^ or ΔZBD-RNAP^549^ was present on DNA.

### BCM growth assays

RapA KO strain (F-, *ΔhepA769::kan*, *Δ(araD-araB)567*, *ΔlacZ4787*(::rrnB-3), *λ^-^*, *rph-1*, *Δ(rhaD-rhaB)568*, *hsdR514* – Keio collection, National BioResource Project entry JW0058) and WT parental strain (E. Coli BW25113 - National BioResource Project entry ME9062) were both separately transformed with plasmids pBAD18 (empty vector) ^70^ or pBAD18*rnhA* (RNase HI) ^47^ and grown overnight on LB-agar plates with 100 μg/mL ampicillin. Colonies were picked and grown overnight in LB with 100 μg/mL ampicillin, then back diluted to an OD_600 nm_ of 0.01 in LB. L-arabinose was added to a final concentration of 0.05% (w/v) for induction. Culture solutions (50 μL) were dispensed into dark well tissue culture plates (Greiner Bio-One CELLSTAR, 384-well) along with BCM (when used, from a 2.5 g/L stock in DMSO; Santa Cruz Biotechnology, CAS 38129-37-2) using an HP D300e Digital Dispenser (Tecan). Cells were incubated at 37°C and OD_600 nm_ values were recorded using a plate reader (TECAN). The minimum inhibitory concentration (MIC) for BCM was determined to be 37.5 mg/L.

### NaCl growth assays

RapA KO strain (F-, *ΔhepA769::kan*, *Δ(araD-araB)567*, *ΔlacZ4787*(::rrnB-3), *λ^-^*, *rph-1*, *Δ(rhaD-rhaB)568*, *hsdR514* – Keio collection, National BioResource Project entry JW0058) and WT parental strain (E. Coli BW25113 - National BioResource Project entry ME9062) both separately transformed with plasmids pBAD18 (empty vector) and pBAD18rnhA and grown overnight on 100 ug/ml Ampicillin LB- agar plates. Colonies were picked and grown overnight in Luria broth with 100 ug/ml Ampicillin. Culture concentrations were standardized according to OD_600 nm_ measurements and were diluted in a 1/10 dilution series, using LB combined with a final concentration of 0.05% arabinose. 2 ml samples from the 1/10 dilution series of all four culture strains were plated onto separate quadrants of LB-agar plates composed of LB-agar, 0.9 M NaCl, and 0.05% arabinose. Plates were incubated at 37°C for 48 hours and then photographed.

## Data availability

All unique/stable reagents generated in this study are available without restriction from the lead contact, Seth A. Darst. The cryo-EM density maps and atomic coordinates have been deposited in the EMDataBank and Protein Data Bank as follows: rPTCc (EMD-40930, PDB 8T00), rPTCi (EMD-40931, PDB 8T02), rPTCo (EMD- 40922, PDB 8SZW), (EMD-26645, 7UOB), RapA^†^-PTC (EMD-40943, PDB 80TL). The atomic models used for initial model building and analysis are available from the Protein Data Bank under the accession codes 6GH6, 6ALH, and 7M8E.

## Acknowledgments

We thank C. Gross, and members of the Darst-Campbell and Gelles Laboratories for helpful discussions, M. Drolet for pBAD18 and pBAD18*rnhA*, and M. Ebrahim, J. Sotiris, and H. Ng at The Rockefeller University Evelyn Gruss Lipper Cryo-electron Microscopy Resource Center for help with cryo-EM data collection and analysis. Some of the work reported here was conducted at the Simons Electron Microscopy Center (SEMC) and the National Resource for Automated Molecular Microscopy (NRAMM) located at the New York Structural Biology Center, supported by grants from the NIH National Institute of General Medical Sciences (P41 GM103310), NYSTAR, the Simons Foundation (SF349247), the NIH Common Fund Transformative High Resolution Cryo-Electron Microscopy program (U24 GM129539) and NY State Assembly Majority. This work was supported by NIH grants P41 GM109824 and P41 GM103314 to B.T.C, R01 GM38330 to R.L., R01 GM081648 to J.G., and R35 GM118130 to S.A.D.

## Author contributions

Conceptualization; J.B., K.I., J.G., E.A.C., S.A.D. Cloning, protein purification, biochemistry; J.B., K.I., R.M. Mass spectrometry; P.D.B.O. Cryo-EM specimen preparation, data collection, and processing: J.B., L.P. Model building and structural analysis: J.B., L.P., E.A.C., S.A.D. Single-molecule experiments; K.I., J.G. Cell growth assays; J.B., B.B. Funding acquisition and supervision: B.T.C., R.L., J.G., E.A.C., S.A.D. Manuscript first draft: J.B., S.A.D. All authors contributed to finalizing the written manuscript.

## Competing interests

The authors declare there are no competing interests.

Supplementary information is available for this paper.

Correspondence and requests for materials should be addressed to Seth A. Darst.

## Extended Data Figures

**Extended Data Fig 1.**
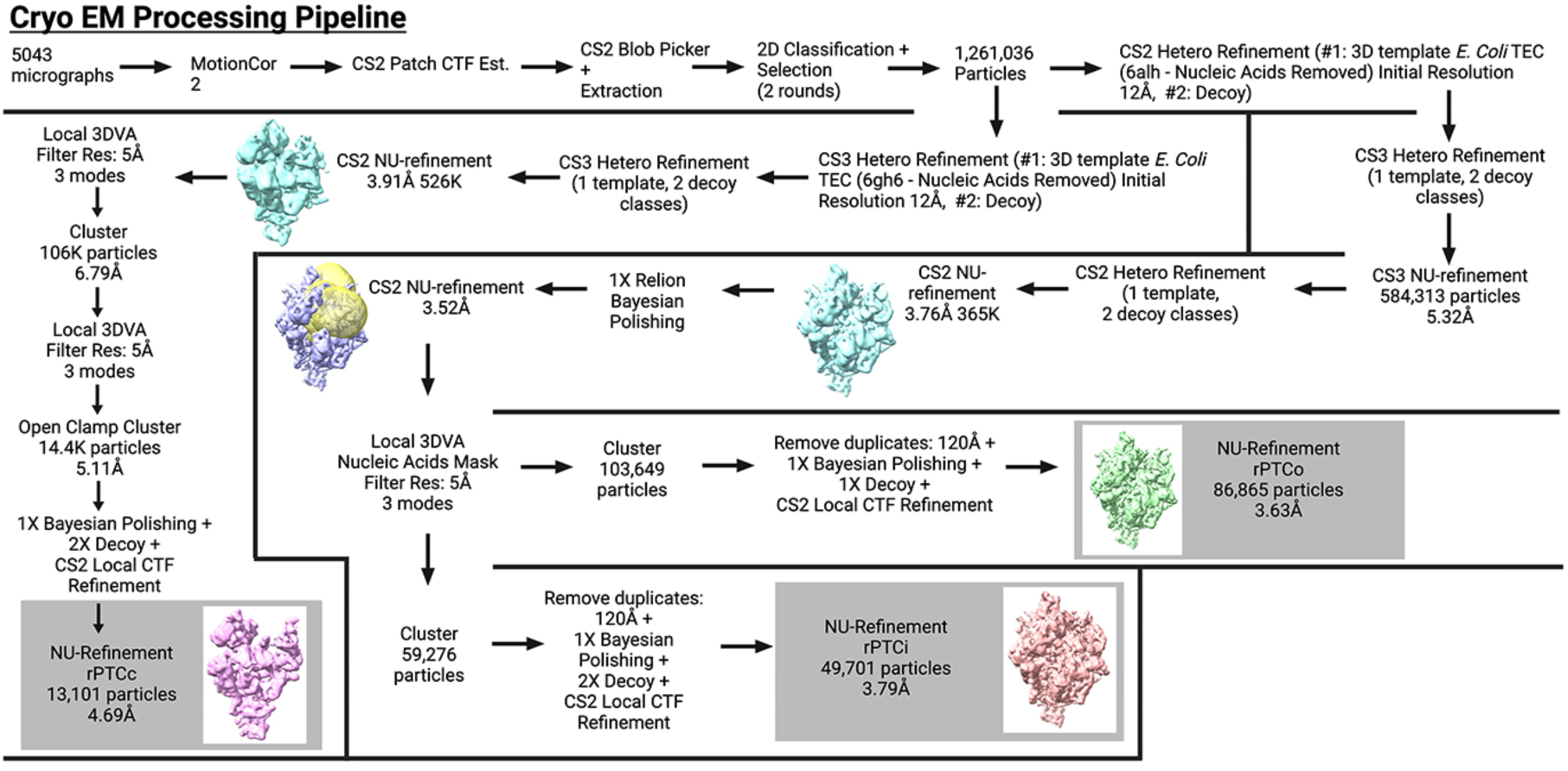
Cryo-EM processing pipeline for rPTC structures (rPTCo, rPTCi, rPTCc).

**Extended Data Fig 2.**
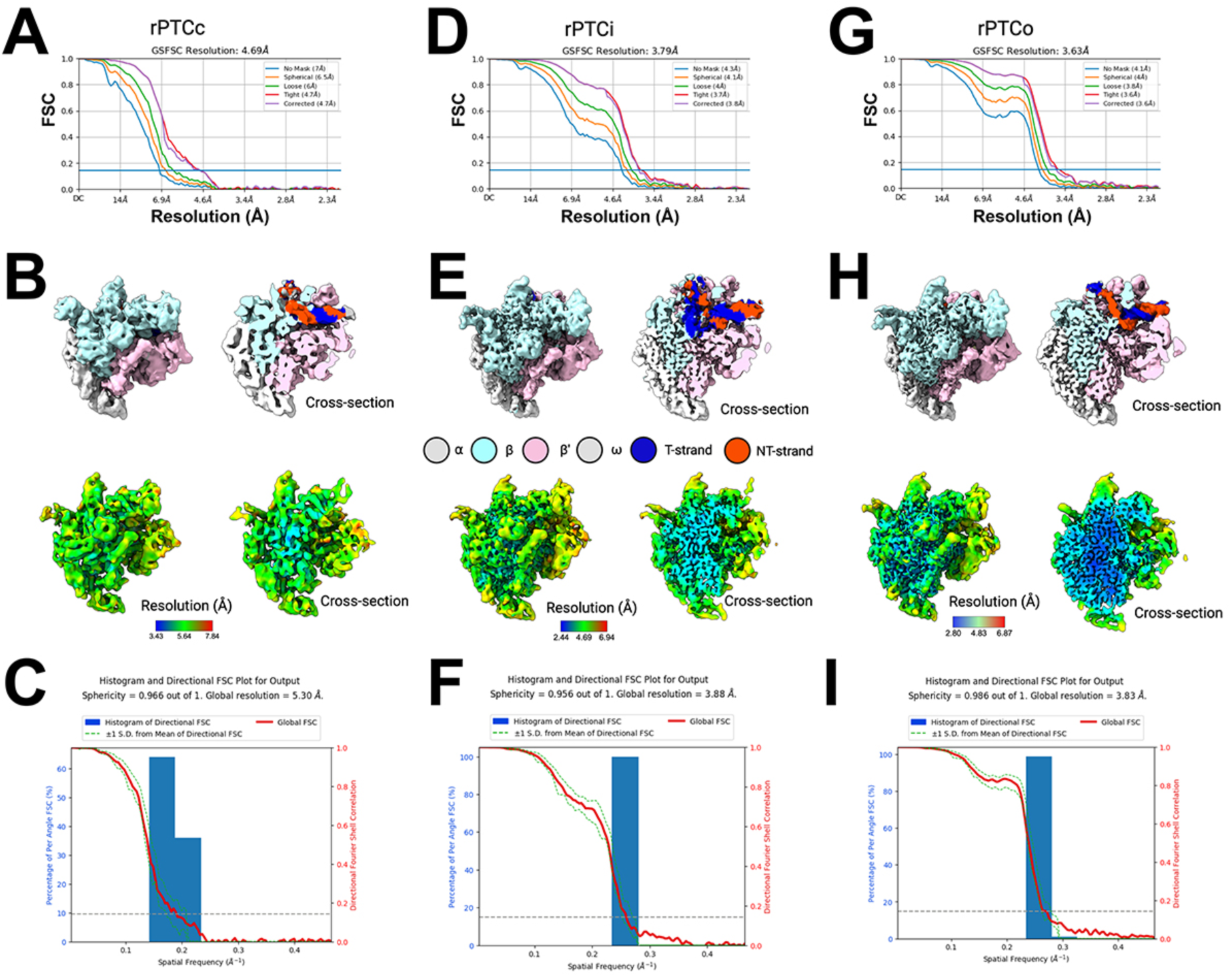
Cryo-EM of rPTCo, rPTCi, and rPTCc. **A, B, C.** rPTCc class: Gold standard FSC calculations for cryo-EM density map (**A**), cryo-EM density map and cross section colored according to key (top of **B**), cryo-EM density map colored according to local resolution (bottom of **B**) ^71^, 3DFSC and sphericity of density map (**C**) ^82^. **D, E, F.** rPTCi class: Gold standard FSC calculations for cryo-EM density map (**D**), cryo-EM density map and cross section colored according to key (top of **E**), cryo-EM density map colored according to local resolution (bottom of **E**) ^71^, 3DFSC and sphericity of density map (**F**) ^82^. **G, H, I.** rPTCo class: Gold standard FSC calculations for cryo-EM density map (**G**), cryo-EM density map and cross section colored according to key (top of **H**), cryo-EM density map colored according to local resolution (bottom of **H**) ^71^, 3DFSC and sphericity of density map (**I**) ^82^.

**Extended Data Fig 3.**
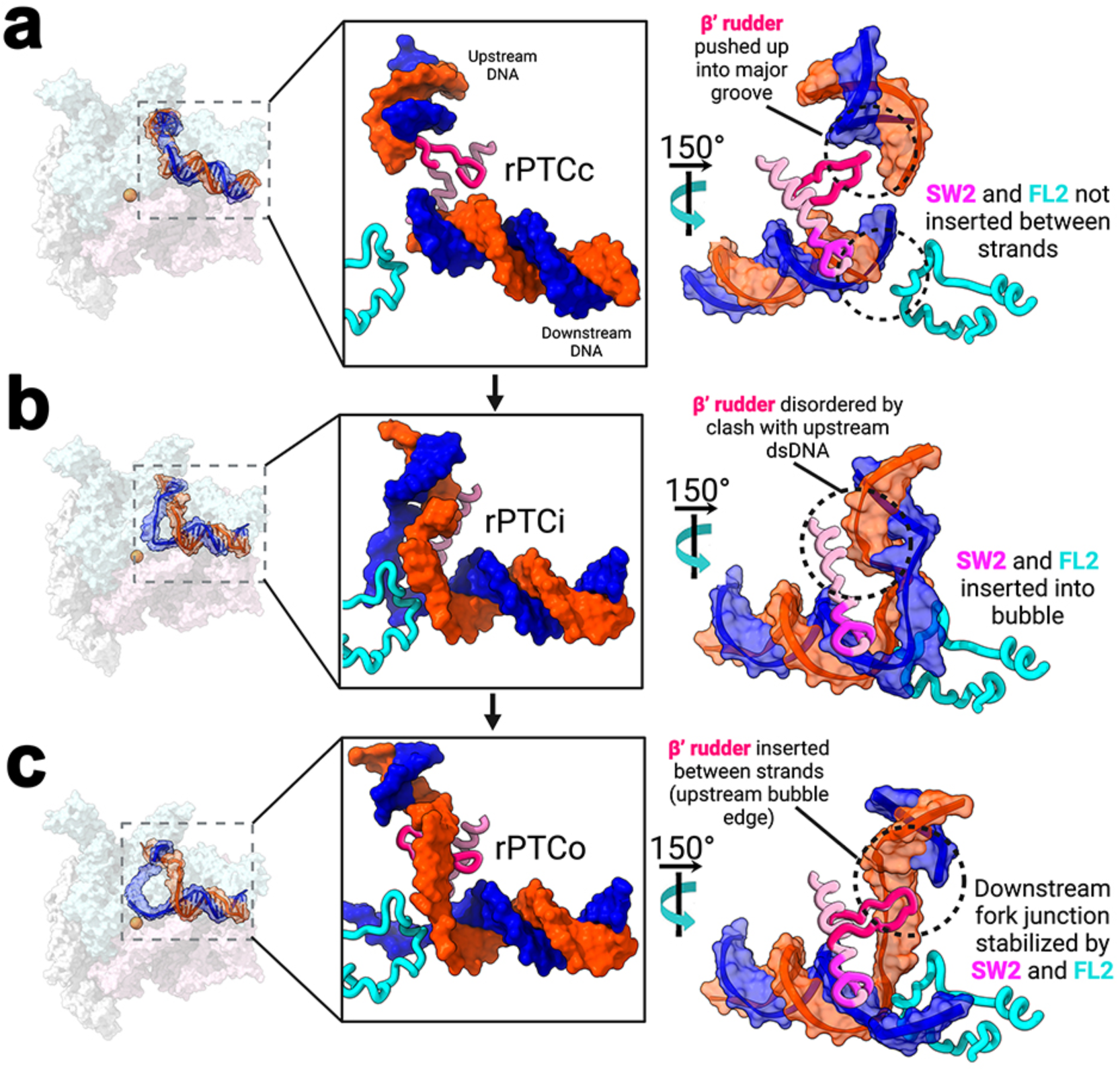
The role of RNAP structural elements in DNA melting. **A.-C.** (*left*) View of the rPTC structures (same as Figs. 1c-e). The boxed regions are magnified on the right. (*right*) Magnified views of boxed region; only the DNA, βFork-loop2 (FL2), β’rudder, and β’Switch2 (Sw2) are shown. DNA is shown as a backbone cartoon with a transparent molecular surface (t-strand, blue; nt-strand, orange). The protein elements are shown as backbone cartoons (β, cyan; β’, pink). **A.** rPTCc; the clamp is open 24°, resulting in a large separation between FL2 and Sw2. **B.** rPTCi; the clamp closes, closing the gap between FL2 and Sw2, nucleating a ∼5 nt bubble in the DNA. The β’rudder is completely disordered. **C.** rPTCo; the bubble propagates in the upstream direction to ∼7-8 nt, creating room for the β’rudder.

**Extended Data Fig. 4.**
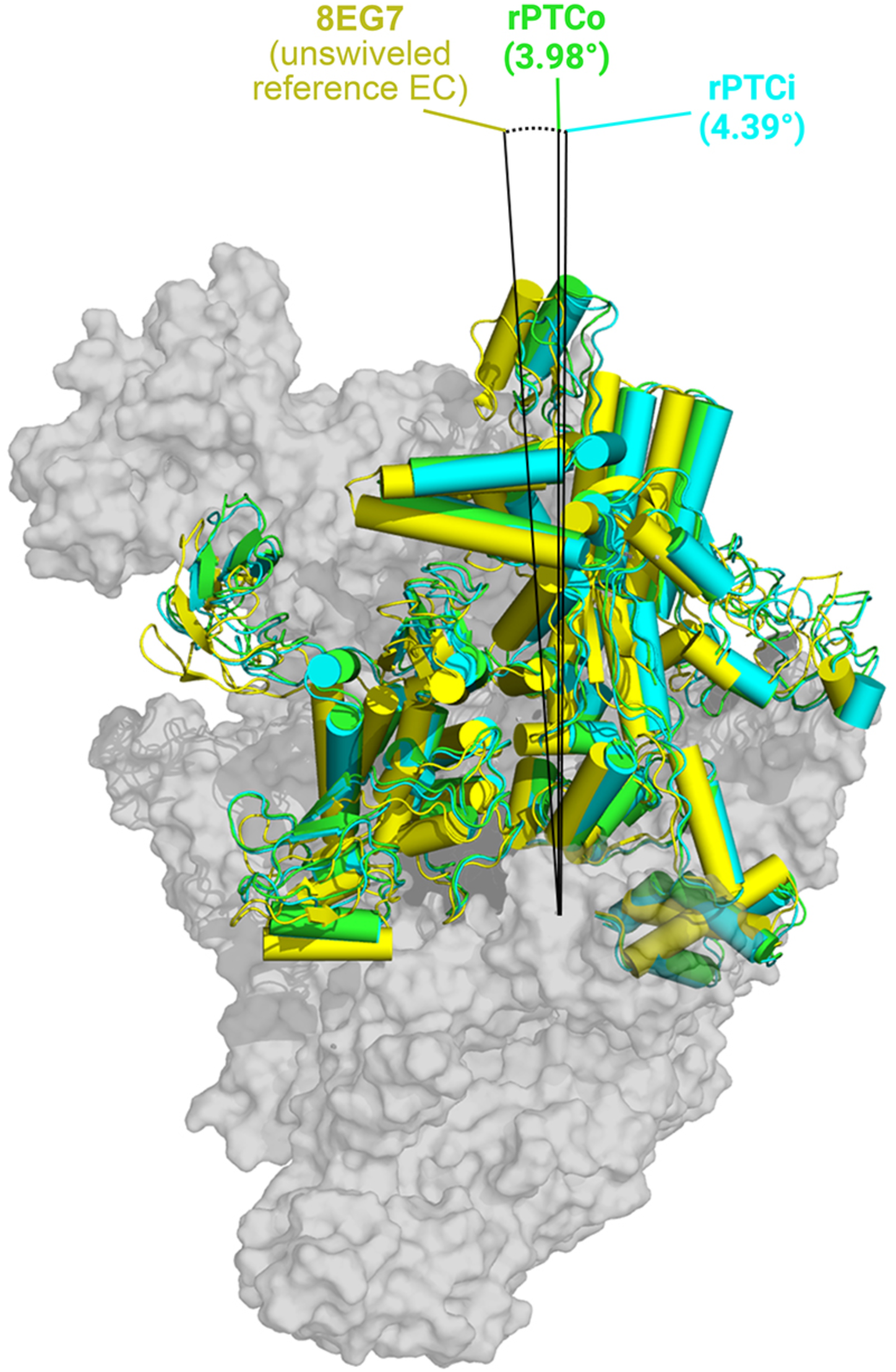
Swiveling in rPTCi and rPTCo.

**Extended Data Fig. 5.**
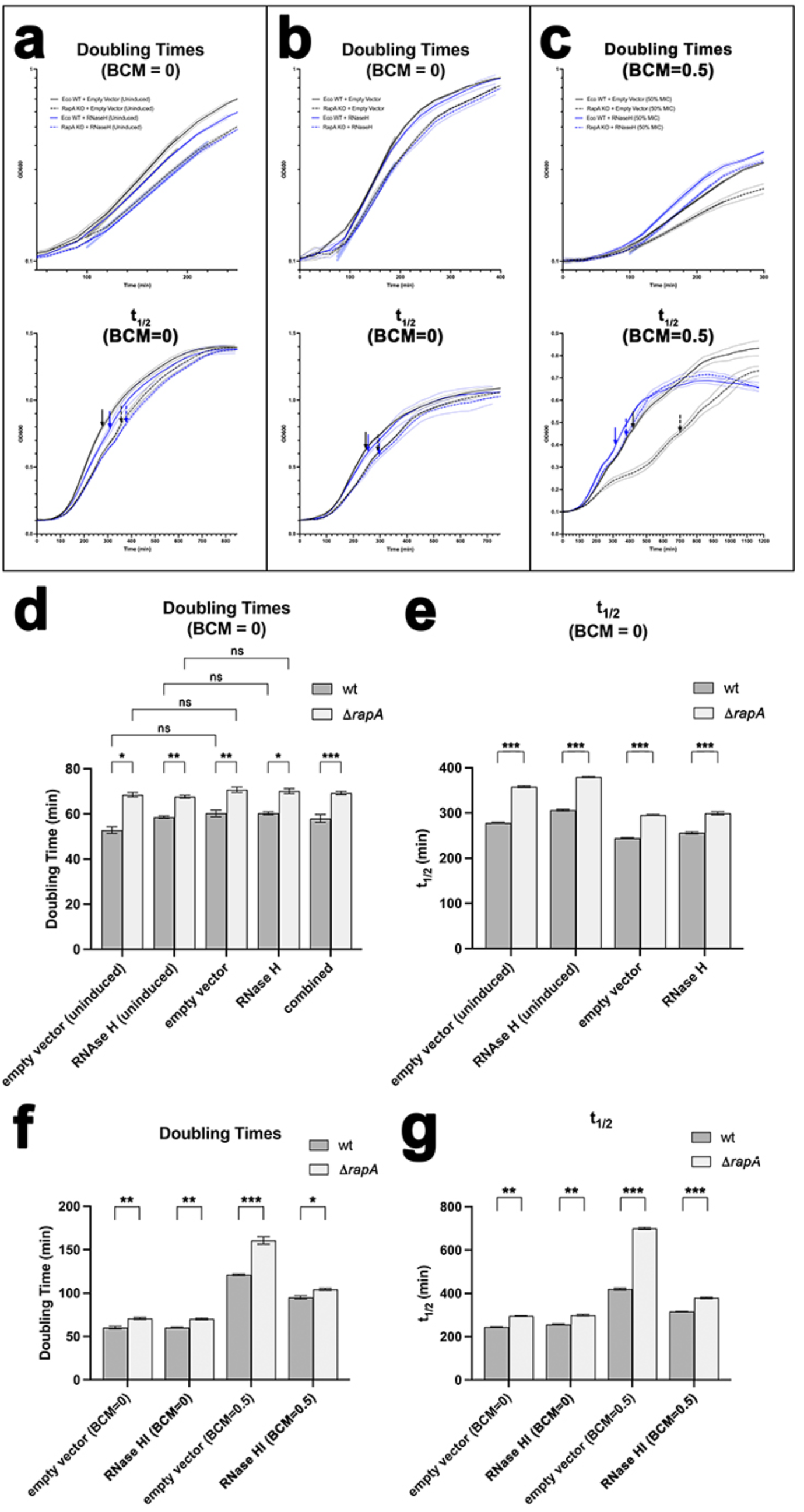
Growth analysis of wt and Δ*rapA Eco*. **a-c.** Growth curves (OD_600 nm_). The solid lines plot the average of three replicates, the thin lines above and below show the 95% confidence limit. (*top panel*) Semi-log plots of the growth curves during log-phase growth. The thick, transparent lines show the exponential fit used to calculate the doubling time. (*bottom panel*) Linear scale showing the full growth curves. The thick arrows denote the t_1/2_. **a.** BCM=0 and in the absence of arabinose [uninduced pBAD18 (empty vector) or pBAD18*rnhA* (RNase H)]. **b.** BCM=0 but with arabinose induction (0.05% w/v) of pBAD18 (empty vector) or pBAD18*rnhA* (RNase H). **c.** BCM=0.5X MIC (MIC = 37.5 mg/L) and with arabinose induction (0.05% w/v) of pBAD18 (empty vector) or pBAD18*rnhA* (RNase H). **d.-e.** Histograms showing growth parameters [double times (**d**) and t_1/2_ (**e**)) for wt and Δ*rapA Eco* cells carrying pBAD18 (empty vector) ^70^ or pBAD18*rnhA* (RNase HI) ^47^, all without BCM (BCM=0). In **d.**, combined is the average for all the measurements. Error bars denote standard error (N=3 or 4 measurements). Statistical significance of differences between samples was determined using unpaired, two-tailed *t*-test (ns, *P* > 0.05; *, *P* ≤ 0.05; **, *P* ≤ 0.01; ***, *P* ≤ 0.001). **f.-g.** Histograms showing growth parameters (doubling times and t_1/2_;) for wt and Δ*rapA Eco* cells carrying pBAD18 (empty vector) ^70^ or pBAD18*rnhA* (RNase HI) ^47^ without (BCM=0) or with 0.5X MIC BCM (BCM=0.5). Error bars denote standard error (≥ N=3 measurements). Statistical significance of differences between samples was determined using an unpaired, two-tailed *t*-test (ns, *P* > 0.05; *, *P* ≤ 0.05; **, *P* ≤ 0.01; ***, *P* ≤ 0.001). **f.** Doubling times. **g.** t_1/2_.

**Extended Data Fig 6.**
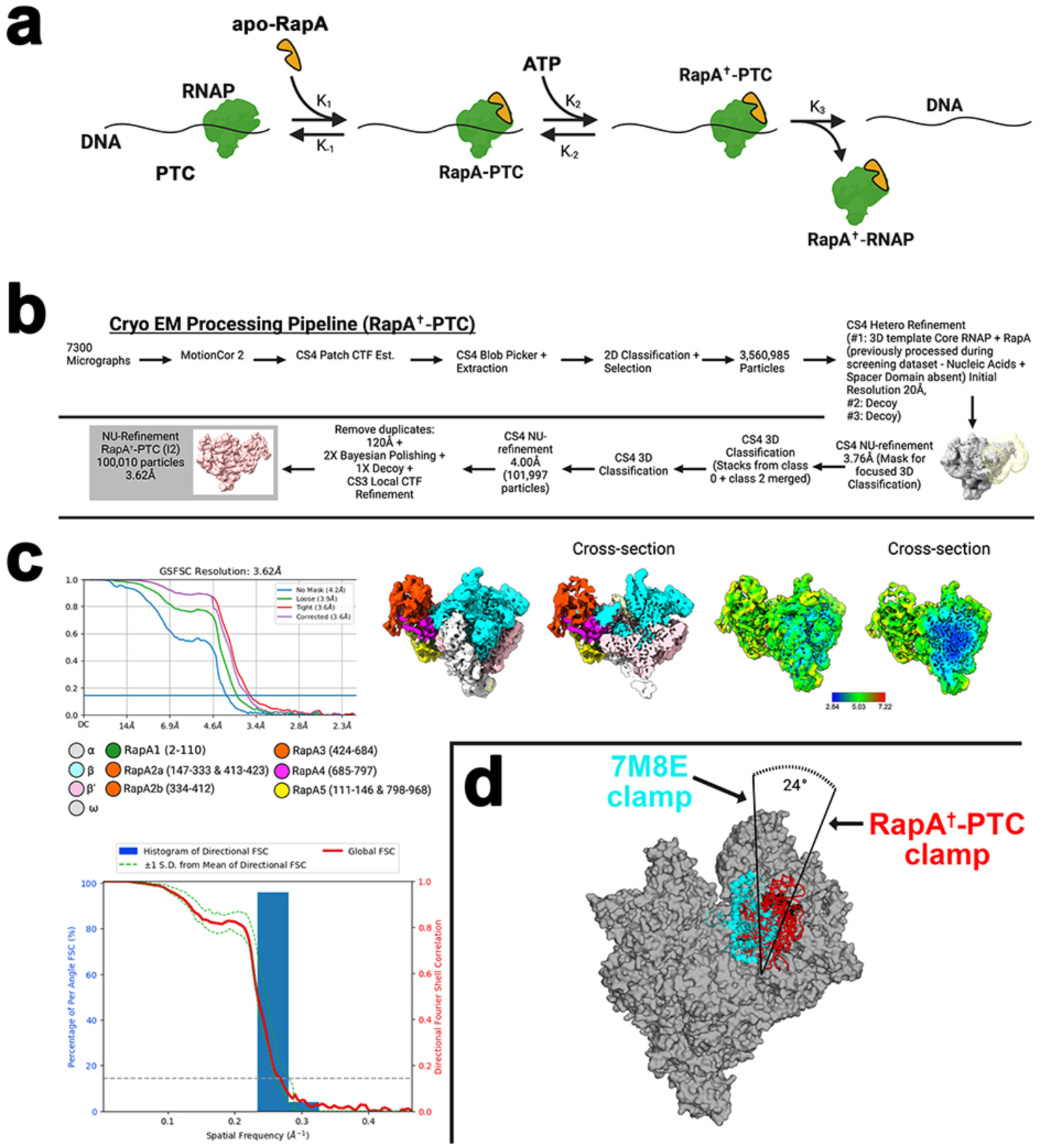
RapA^†^-PTC. **a.** Hypothesized mechanism for the disruption of the PTC by RapA - adapted from ^7^. **b.** Cryo-EM processing pipeline for RapA^†^-PTC class. **c.** Gold standard FSC calculations for the cryo-EM density map (upper left), cryo-EM density map and cross section colored according to key (upper middle), cryo-EM density map colored according to local resolution (upper right) ^71^, 3DFSC and sphericity of density map (lower) ^82^. **d.** Range of RNAP clamp movement observed in the transition between RapA-PTC stand-in (PDB: 7M8E) and observed RapA^†^-PTC. Clamp opens approximately 24° upon RapA-PTC conformation change driven by ATP binding.

**Extended Data Fig. 7.**
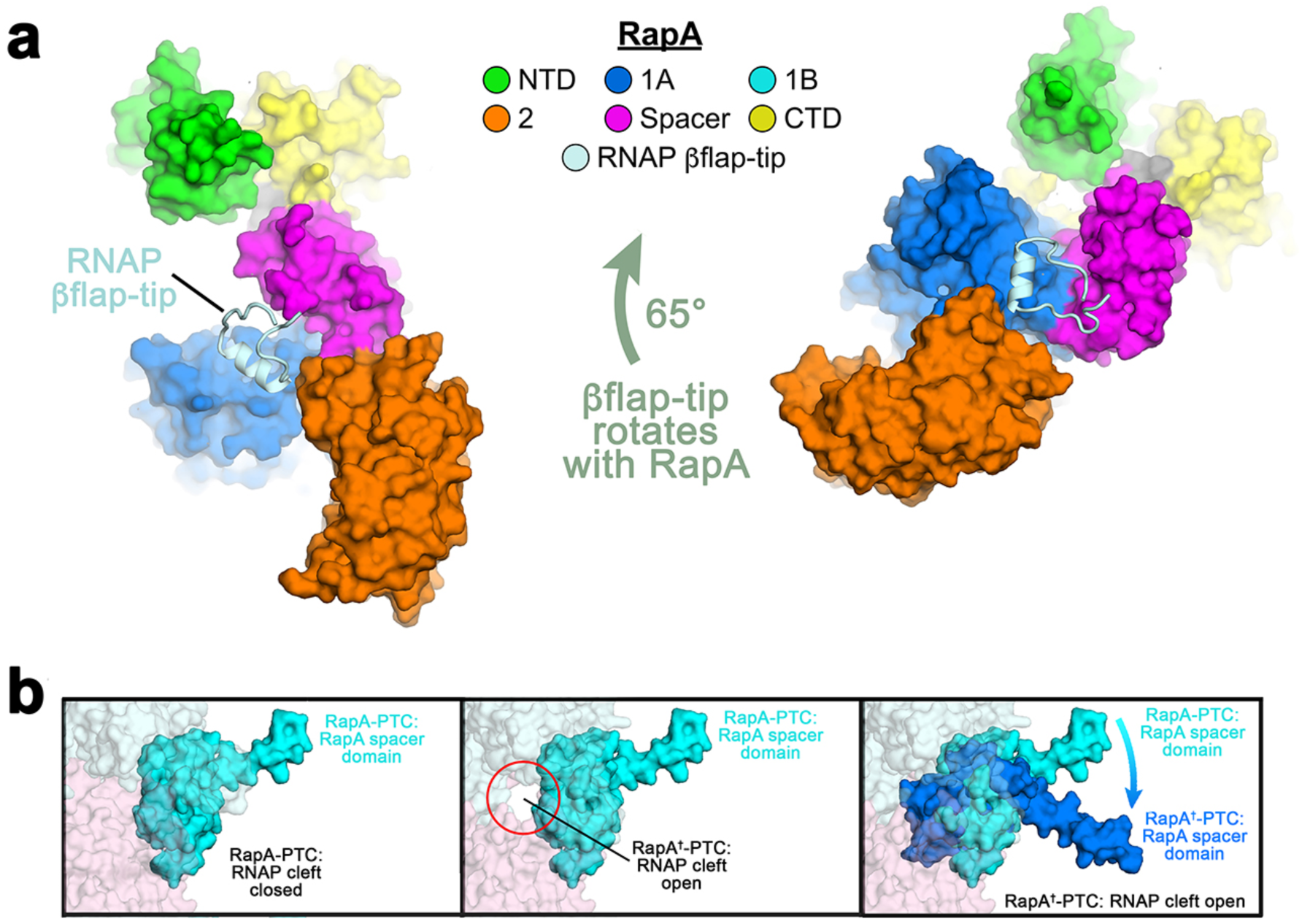
Details of RapA^†^-PTC structural rearrangements. **a.** The overall rotation of RapA^†^ with respect to the RNAP (65°, see Fig. 4c) is acommodated by flexibility of the RNAP βflap-tip, which maintains its contacts with RapA but also rotates with respect to the rest of the RNAP. **b.** The RNAP cleft opens as the RNAP clamp is pulled open by the RapA^†^ structural rearrangements (transition from left to middle panel; also see Fig. 4d). The RapA^†^ spacer domain (cyan to dark blue transition) wedges into the open RNAP cleft (transition from middle to right panel).

## SUPPLEMENTAL INFORMATION

**Supplementary Table 1.** Cryo-EM data collection, refinement, and validation statistics for RapA^†^-PTC and rPTC classes.

| Dataset | rPTC |  |  | RapA<sup>†</sup>-PTC |
| --- | --- | --- | --- | --- |
| Data collection and processing |  |  |  |  |
| Microscope | FEI Titan Krios |  |  | FEI Titan Krios |
| Voltage (kV) | 300 |  |  | 300 |
| Detector | Gatan K3 |  |  | Gatan K3 |
| Electron exposure (e-/Å<sup>2</sup>) | 55.9 |  |  | 52.1 |
| Defocus range (μm) | -0.25 to -4.2 |  |  | -0.8 to -2.5 |
| Data collection mode | Counting Mode |  |  | Counting Mode |
| Pixel size (Å) | 1.076 |  |  | 1.076 |
| Symmetry imposed | C1 |  |  | C1 |
| Initial particle images (no.) | 1,261,036 |  |  | 3,560,985 |
| Refinement |  |  |  |  |
| Structure | rPTCc | rPTCi | rPTCo | RapA<sup>†</sup>-PTC |
| EMDB | EMD-40930 | EMD-40931 | EMD-40922 | EMD-40943 |
| PDB | 8T00 | 8T02 | 8SZW | 8T0L |
| Final particle images (no.) | 13,101 | 49,701 | 86,865 | 100,010 |
| Map resolution (Å) - FSC threshold 0.143 | 4.7 | 3.8 | 3.6 | 3.6 |
| Map resolution range (Å) | 3.4 - 7.8 | 2.4 - 6.9 | 2.8 - 6.9 | 2.8 - 7.2 |
| Initial model used (PDB code) | 6GH6 | 6ALH | 6ALH | Ab-initio via screening dataset |
| Map sharpening B factor (Å<sup>2</sup>) | -126.0 | -108.9 | -114.5 | -117.5 |
| Model composition |  |  |  |  |
| Non-hydrogen atoms | 50,247 | 51,102 | 51,717 | 65,326 |
| Protein residues | 3,093 | 3,146 | 3,183 | 4,152 |
| Nucleic acid residues (DNA) | 52 | 51 | 52 | - |
| Ligands | 2 Zn<sup>2+</sup>, 1 Mg<sup>2+</sup> | 2 Zn<sup>2+</sup>, 1 Mg<sup>2+</sup> | 2 Zn<sup>2+</sup>, 1 Mg<sup>2+</sup> | 2 Zn<sup>2+</sup>, 1 Mg<sup>2+</sup>, 1 ADP-AlF<sub>3</sub> |
| B factors (Å<sup>2</sup>) |  |  |  |  |
| Protein | 289.6 | 109.2 | 69.94 | 166.2 |
| Nucleic acid | 515.0 | 247.9 | 134.2 | - |
| Ligands | 382.0 | 139.8 | 83.67 | 233.2 |
| R.m.s. deviations |  |  |  |  |
| Bond lengths (Å) | 0.003 | 0.004 | 0.003 | 0.005 |
| Bond angles (°) | 0.662 | 0.643 | 0.63 | 0.668 |
| Validation |  |  |  |  |
| MolProbity score | 1.82 | 1.99 | 1.83 | 2.62 |
| Clashscore | 7.07 | 9.67 | 7.68 | 9.74 |
| Poor rotamers (%) | 0 | 0.22 | 0.22 | 5.59 |
| Ramachandran plot |  |  |  |  |
| Favored (%) | 93.49 | 92.06 | 93.93 | 90.2 |
| Allowed (%) | 6.48 | 7.91 | 5.97 | 9.68 |
| Disallowed (%) | 0.03 | 0.03 | 0.09 | 0.12 |

**Supplementary Table 2.** RapA domains.

| *Eco* RapA residues | domain | color |
| --- | --- | --- |
| 1-110 | NTD | green |
| 111-146 | Linker1 |  |
| 147-333/412-423 | RecA-1A | marine |
| 334-411 | 1B | cyan |
| 424-427 | Linker2 |  |
| 428-683 | RecA-2 | orange |
| 684-712 | Linker3 |  |
| 713-793 | Spacer | magenta |
| 112-142/798-C-term | CTD | yellow |

**Supplementary Table 3.** Comparison of rPTCc, RapA^†^-PTC, and 8AC2.

|  | rPTCc | RapA<sup>†</sup>-PTC |
| --- | --- | --- |
| rPTCc |  |  |
| RapA<sup>†</sup>-PTC | 0.983 Å<sup>a</sup><br>(2,858 Cαs) | |
| 8AC2 | 0.994 Å<br>(2,748 Cαs) | 1.257 Å<br>(2,937 Cαs) |

<sup>a</sup>Top number is the root-mean-square-deviation (rmsd). Below in parenthesis is the number of $\alpha$ -carbons compared in the rmsd calculation.

## Supplementary Videos

**Supplementary Video 1. RapA conformational changes induced by nucleotide binding.** The video illustrates the RapA conformational changes induced by nucleotide (ADP-AlF_3_) binding. The video starts by showing the EC from an apo-RapA-EC structure (7M8E ^21^), then showing apo-RapA. The video then focuses on RapA and highlights the conformational transitions between apo-RapA and RapA(ADP-AlF_3_).

**Supplementary Video 2. RNAP conformational changes mediated by RapA.** The video illustrates the RNAP conformational changes mediated by the RapA conformational changes. The video starts showing the EC from an apo-RapA-EC structure (7M8E ^21^), with the RNAP clamp and β’ZBD highlighted. Apo-RapA is then shown. The video then splits into two screens, each showing a unique orientation of the RapA-EC structure, which transitions back and forth from the RapA-EC to the RapA^†^-EC. The left screen illustrates the large motions of RapA with respect to the RNAP. The right screen illustrates how the RapA conformational changes cause RNAP clamp opening.

## Notes

### Competing Interest Statement

The authors have declared no competing interest.

